# A paleogenomic and paleoclimatic approach to reconstruct historical responses of coral holobionts to anthropogenic change

**DOI:** 10.1101/2024.09.02.610915

**Authors:** Raúl A. González-Pech, Kelsey Dyez, Luis Lizcano-Sandoval, Raphael A Eisenhofer-Philipona, Stephanie Marciniak, Colin Howe, Sofia Roitman, Ángela M. Marulanda-Gómez, Alberto Rodriguez-Ramirez, Natalie A. Falta, Benjamin Knowles, George H. Perry, Laura S. Weyrich, Mateo López-Victoria, Julia E. Cole, Mónica Medina

## Abstract

Coral reefs are declining worldwide due to anthropogenic environmental change. The foundation and health of these ecosystems rely on the harmonious functioning of all members of coral holobionts, i.e., cnidarian host, symbiotic microalgae, and associated microbiome. Coral stress responses often involve shifts in the taxonomic identity of their symbionts and microbiomes. Tracing back changes in coral holobiont composition over prolonged time periods can help us reconstruct health history of reefs and gain a better understanding of coral response to current stressors. Here, we focused on a major Caribbean reef-builder coral, *Orbicella faveolata*, from Varadero Reef, Colombia. This reef has undergone extensive freshwater sediment discharge and pollution for decades as result of urbanization. We show, for the first time, that paleogenomic and paleoclimatic approaches can be combined to reconstruct historical coral holobiont dynamics potentially associated with anthropogenic disturbances.

## Introduction

The past century has witnessed environmental disturbances with unprecedented rates of change in human history; these changes are driving profound losses of biodiversity worldwide^1–3^. Coral reefs are among the ecosystems with the most biodiversity and highest productivity in the planet^4,5^, but they are also among the most impacted by anthropogenically induced climate change^6^. Increasingly frequent and severe marine heat waves spanning the global tropics are leading to mass bleaching events (in which corals lose their symbionts due to oxidative stress^7^) followed by widespread mortality^8,9^. Global temperature stress is exacerbated by local impacts, such as increased sedimentation^10^, overfishing^11^, nutrification^12^, other coastal threats^13^, and their interactions^14^. These threats cause major biodiversity shifts^15^ that trickle down to microbial scales^16^, which in turn alter coral reef functioning and critically undermine ecosystem services for hundreds of millions of people^17,18^.

Paradoxically, coral reefs have been thriving in the nutrient-poor (i.e., oligotrophic), shallow waters of tropical oceans for the past 200 million years. This is thanks to the obligate endosymbioses between reef-building corals and photosynthetic unicellular dinoflagellates in the family Symbiodiniaceae^19,20^. The nutritional exchange taking place in these symbioses provides the metabolic energy that enable reef-building corals to deposit calcium carbonate at rates exceeding those of erosion^21,22^, resulting in the formation of tridimensional reef structures^23^. Coral calcification is strongly dependent on endosymbiont photosynthesis^24^, thus changes in water optical properties associated with local disturbances have negative effects on the capacity of corals to maintain functional integrity in impacted reefs^25^. Long-term declines in water quality on reefs have been mainly linked to increased sediment, nutrient, and pollutant loads delivered from adjacent river catchments^26,27^.

The term ‘coral holobiont’ describes the combination of animal host, algal symbionts, and associated prokaryotic, protistan, and viral assemblages that collectively interact with local environmental conditions to determine coral phenotype^28–31^. Bacteria and archaea might be important for carbon fixation and nitrogen metabolism in the holobiont, as well as in providing vitamins and amino acids to the eukaryotic partners^32^. Bacterial partners supplement the metabolic demands of coral hosts and the dinoflagellate symbionts while coping with heat stress^33^. Endolithic communities (i.e., bacteria and algae living within the skeleton) also appear to play important roles in coral holobiont health^34,35^. Viruses might affect microbial dynamics, bleaching, and disease, and even play a role in reef biogeochemical cycling^36^. Therefore, the functioning of a coral holobiont is contingent on the interactions among all its members and the ongoing environmental conditions at a certain timepoint.

Corals preserve a record of their physical and chemical environment in their skeletal geochemistry and growth banding. Because some massive coral species can live for centuries, they can reveal detailed histories of environmental fluctuations that characterize disturbances on reefs over the Anthropocene. Coral skeletons therefore serve as natural archives storing detailed records of past environments^37–39^. Skeletal growth and chemical composition can be quantified to indicate changes in calcification, extension, density, and geochemical composition, which may reveal environmental changes in temperature, salinity, alkalinity, sedimentation, nutrient availability, and sea level^26,40–47^. Combined analyses of coral growth and calcification rates (via sclerochronology) with geochemical records (stable isotopes and trace elements) provide thus powerful tools to reconstruct historical environmental reef stressors associated with human activities, such as changes in sea surface temperature (SST) and salinity, ocean acidification, runoffs, and pollution. Because some massive coral species can live for centuries, skeletons from these corals can reveal detailed histories of environmental fluctuations that characterize disturbances on reefs over the Anthropocene. The most recent analyses of 400 year-old coral skeleton cores from the Australian Great Barrier Reef^48^ and a 600 year-old core from Fiji^49^ revealed evidence of human-induced warming leading to the rapid increase of mass bleaching events.

The implementation of historical and ancient DNA (aDNA) techniques can be used to explore new chapters in the skeletal archives of corals. A previous study combined aDNA techniques with *ITS2*-amplicon sequencing to search for shifts of dinoflagellate symbionts in soft coral specimens spanning 172 years from a museum collection^50^. The study detected the same symbiont type in historical and modern samples across six host species, thus revealing strong stability of coral-dinoflagellate associations over time. A recent investigation came to the same conclusion following a similar approach on freshly sampled skeleton cores from the massive stony coral *Porites lutea*, with no evidence of changes in the dominant symbiont type for about 150 years^51^. This latter study also coined the term “*cora*DNA” to refer to the implementation of aDNA techniques to the study of corals. Another recent study applied *cora*DNA and metagenomic techniques to millennia-old fossils of the coral *Acropora palmata* to gain insights into adaptive trajectories of the coral holobiont^52^. This work could recover DNA from the coral host and microbiome, but none from the dinoflagellate symbionts. Ancient microbiomes were similar to those of extant acroporids, which also suggests long-term stability and specialization of coral microbiomes. However, to our knowledge, no efforts have been undertaken to take advantage of *cora*DNA to understand how environmental stressors and human activities impact coral holobiont members over time, a strategy that could have a major impact on improving predictive models of holobiont resilience, as well as on the more-informed design of synthetic microbial communities for probiotic treatments under different climate change scenarios.

Here, we sampled a coral skeletal core, spanning ∼65 years, from the massive Caribbean reef-builder *Orbicella faveolata* from the Varadero Reef, Colombia– a reef that has experienced a long history of human impacts^53–56^. We used sclerochronology, geochemistry, *cora*DNA, metagenomics, and historical environmental records to show that we can reconstruct past environmental conditions and responses of the coral holobiont to recent environmental changes. For the first time, we recovered historical DNA from coral host, algal symbionts, and microbiome (including viruses) from a single coral skeletal core, and we paired these with paleoclimatic methods to assess the role of external factors. This study shows that a multi-pronged approach, incorporating both traditional and novel techniques, can be leveraged to successfully reconstruct historical responses of coral holobionts to environmental disturbances.

## Results

### coraDNA recovered from coral skeleton core

We used a core sample from a colony of *Orbicella faveolata* in the Varadero Reef, Colombia^57^ (Figure 1). We used annual cycles in Sr/Ca, Ba/Ca, δ^18^O, and luminescence (see *Experimental procedures*), to determine that the coral core spans ∼65 years (1955–2015; Figure 2a). Subsamples roughly corresponding to one year were taken from the core at ∼5-year intervals (see *Experimental Procedures* and Supplementary Figure 1) for coral ancient DNA analysis (*cora*DNA^51^; Supplementary Table 1). Short DNA fragments and cytosine deamination, common types of damage in ancient DNA (aDNA), were detected in the DNA isolated from subsamples with sufficient data and were more pronounced in earlier timepoints (Supplementary Figure 2), indicating that the recovered DNA is endogenous to the coral.

**Figure 1.**
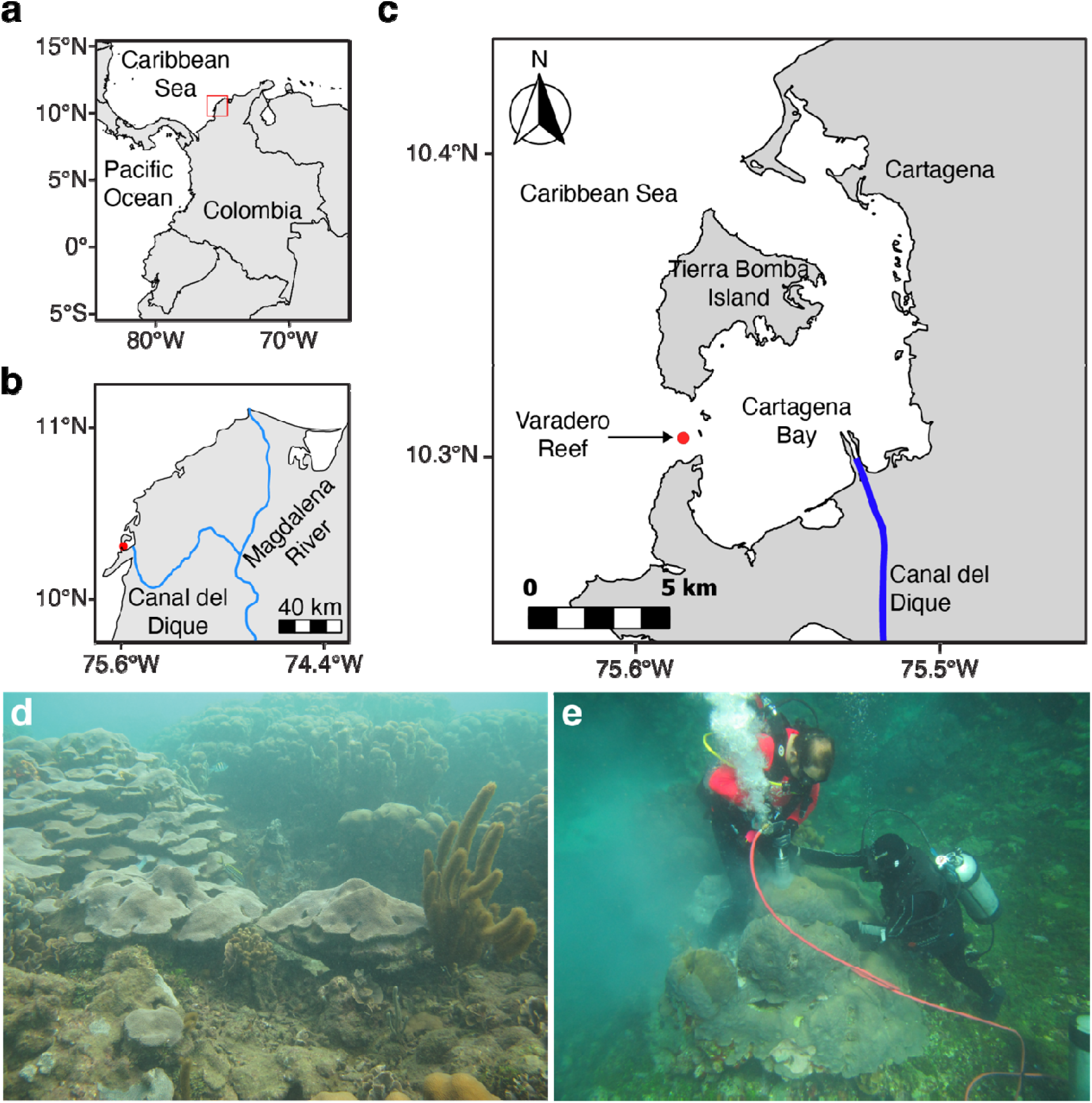
**(a-c)** Location of Varadero Reef in Cartagena Bay relative to the Magdalena River and Canal del Dique. **(d)** Underwater photograph of Varadero Reef showing massive colonies of *Orbicella faveolata* growing at ∼4 m depth; note the turbidity. Photo credit by Melina Rodríguez. **(e)** Underwater photograph showing the sampling procedure by SCUBA using a pneumatic drill. Divers in the photo are authors of this article (Mateo López-Victoria on the left, Alberto Rodriguez-Ramirez on the right). Photo credit by Ángela M. Marulanda-Gómez.

**Figure 2.**
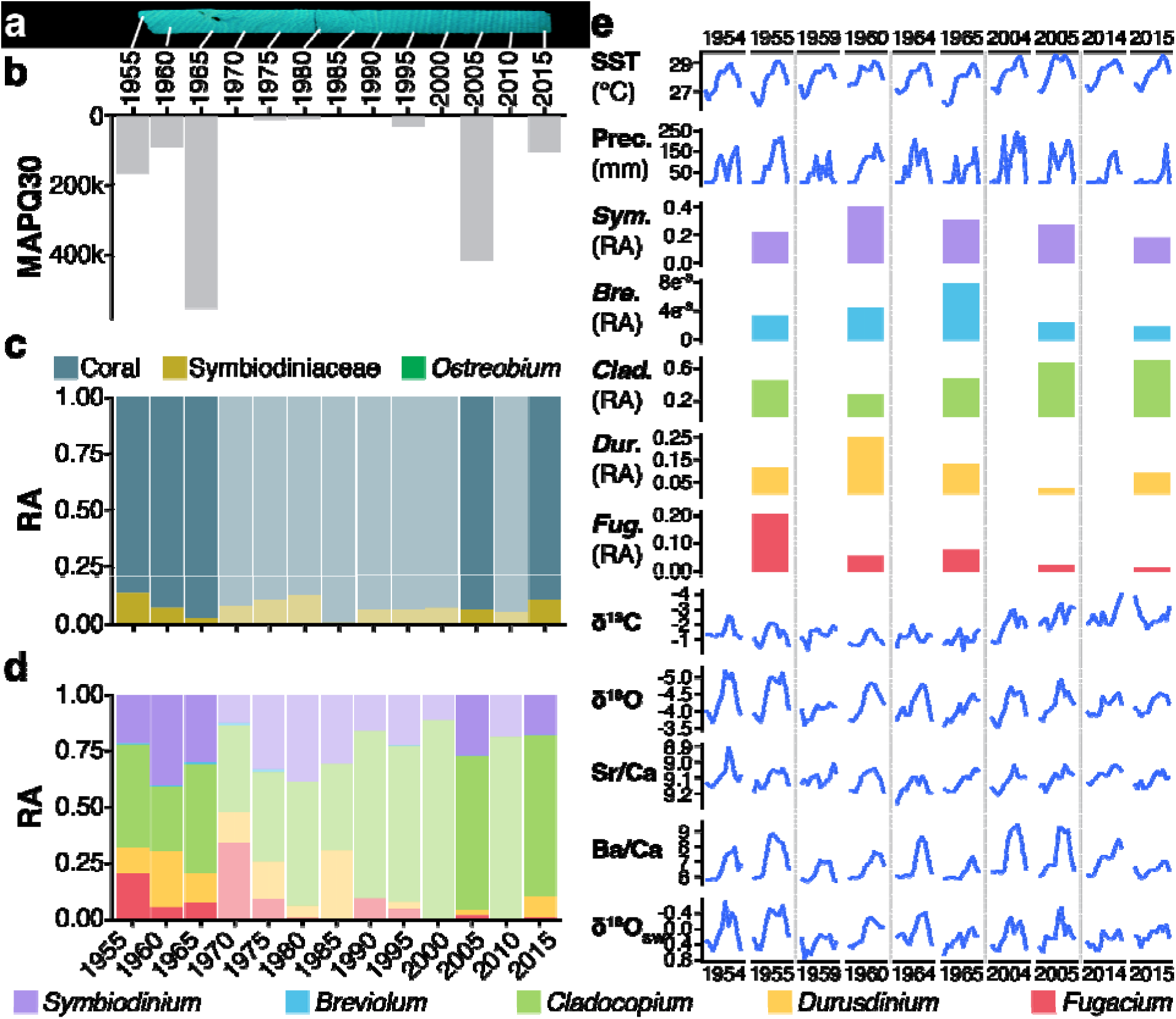
**(a)** Coral skeletal fluorescence (blue-green colors) from the sampled core showing yearly growth bands. **(b)** Number of processed reads that aligned high quality (MAPQ ≥ 30) to genome references of eukaryotic members of the coral holobiont (Supplementary Table 3). **(c)** Relative abundance (RA) of eukaryotic holobiont members (i.e., coral host, Symbiodiniaceae symbionts, and *Ostreobium*), over time based on read alignments in **(b)**. **(d)** Same as **(c)** but only for Symbiodiniaceae and broken down by genus. Transparencies in **(c)** and **(d)** represent subsamples with an initial sequencing depth <5 M reads and sufficient data aligned to the genome references **(b)**. **(e)** Monthly fluctuation of environmental conditions (top), annual relative abundance of Symbiodiniaceae general (middle), and monthly fluctuation of geochemical measurements (bottom) for the years subsampled from the core and the year before. Note that the *y*-axis of δ^13^C, δ^18^O, Sr/Ca, and δ^18^C_SW_ are inverted to reflect their negative correlation with the environmental condition they are proxy of. ‘SST’: sea surface temperature; ‘Prec.’: precipitation; ‘*Sym*.’: *Symbiodinium*; ‘*Bre.*’: *Breviolum*; ‘*Clad.*’: *Cladocopium*; ‘*Dur.*’: *Durusdinium*; ‘*Fug.*’: *Fugacium*.

Stepwise alignment of processed reads to the genome sequences of coral host, Symbiodiniaceae, and *Ostreobium* (see *Experimental Procedures*; Supplementary Table 2), revealed that a small fraction of all reads (8.22%) belonged to eukaryotic members of the holobiont, compared to the larger fraction of prokaryotic reads (47.51%) that aligned to microbial metagenome-assembled genomes (MAGs) generated in this study (see *Experimental Procedures*; Supplementary Figure 3). The number of reads that aligned with high confidence to genome references of coral holobiont members varied drastically among subsamples (Figure 2b). The coral host was most strongly represented among eukaryotic members across the six decades, followed by Symbiodiniaceae by two orders of magnitude (Figure 2c and Supplementary Figure 4). *Ostreobium* was only detected in subsamples ≥5 M reads. Deeper characterization of the unaligned reads (44.27%) showed that these data largely encompassed taxa similar to those of the coral core MAGs as well as other members of the holobiont (Supplementary Figure 3), which is comparable to the fractions of microbial reads estimated by singleM^58^ per subsample (Supplementary Table 3).

### Algal symbiont shuffling follows environmental fluctuations

Relative abundances of Symbiodiniaceae genera showed dominance by the *Symbiodinium* and *Cladocopium* genera with shuffling over time (Figure 2d and Supplementary Figure 4). Overall, these relative abundances correlated better with SST than with precipitation (Supplementary Figure 5 and Supplementary Table 4). Geochemical analyses provide proxies of past environmental conditions (see *Experimental procedures*). Specifically, δ^18^O and Sr/Ca were treated as proxies of SST; δ^13^C and Ba/Ca of water flow from *Canal del Dique*^26^; and δ^18^C_SW_ of both water flow and precipitation^43^. The strongest correlations between Symbiodiniaceae relative abundances and geochemical measurement appears with δ^13^C and Ba/Ca, proxies for water flow from Canal del Dique (Supplementary Figure 6).

Change in relative abundance of each genus is more evident in Figure 2e. The first timepoint, 1955, was the one with the most diverse and even presence of Symbiodiniaceae genera. *Symbiodinium* peaked in 1960, decreasing in relative abundance since, as SST and δ^13^C (i.e., water flow) increased. The opposite pattern is observed for *Cladocopium*, which appears more abundant in recent years (2005 and 2015). *Breviolum* peaked in 1965, which correlates with a drop of the maximum yearly temperature (Supplementary Figure 5). *Durusdinium* and *Fugacium* peaked in 1960 and 1955, respectively.

### Microbial community and function changed over as the environment degraded

We recovered 36 microbial MAGs from the sequenced coral core (Supplementary Table 5), including nine named phyla, ten classes, 15 orders, 14 families, 12 genera, and 13 species. None of the MAGs recovered from the coral core were recovered in those from environmental samples (i.e., seawater and sediment) from Varadero Reef and Rosario Islands (a nearby location; see *Experimental procedures*), suggesting the absence of detectable contamination (Supplementary Figure 7 and Supplementary Table 6). Hierarchical clustering of the MAGs relative abundance grouped subsamples in a chronological fashion (Figure 3a), indicating that the closer in time subsamples were, the more similar their microbial community composition was. Principal Component Analysis (PCA) separated subsamples by time as well (Figure 3b), with two main groups along Principal Component 1 (PC1), one including the most recent timepoints and the other one the earlier ones (Figure 3b). The most represented MAGs for every timepoint can be appreciated from both the heatmap (Figure 3a) and the PCA loading plot (Figure 3c). For example, members of the genus *Marinobacter* were the most abundant in earlier timepoints (1955 and 1960), whereas family Cyclobacteriaceae was the most abundant in 1965, and family Desulfosarcinaceae in more recent timepoints (2005 and 2015). This is supported by their contribution to PC1 (Supplementary Table 7), based on which *Alcanivorax profundimaris* and Thalassobaculaceae also differ substantially when comparing earlier with more recent timepoints. This separation aligns with the decrease in the annual δ^13^C mean (a proxy for water flow increase from Canal del Dique (Figure 3b). However, other environmental variables might be playing a role, as shown by an increase in the annual maximum temperature over the time spanned by the core (Supplementary Figure 8).

**Figure 3.**
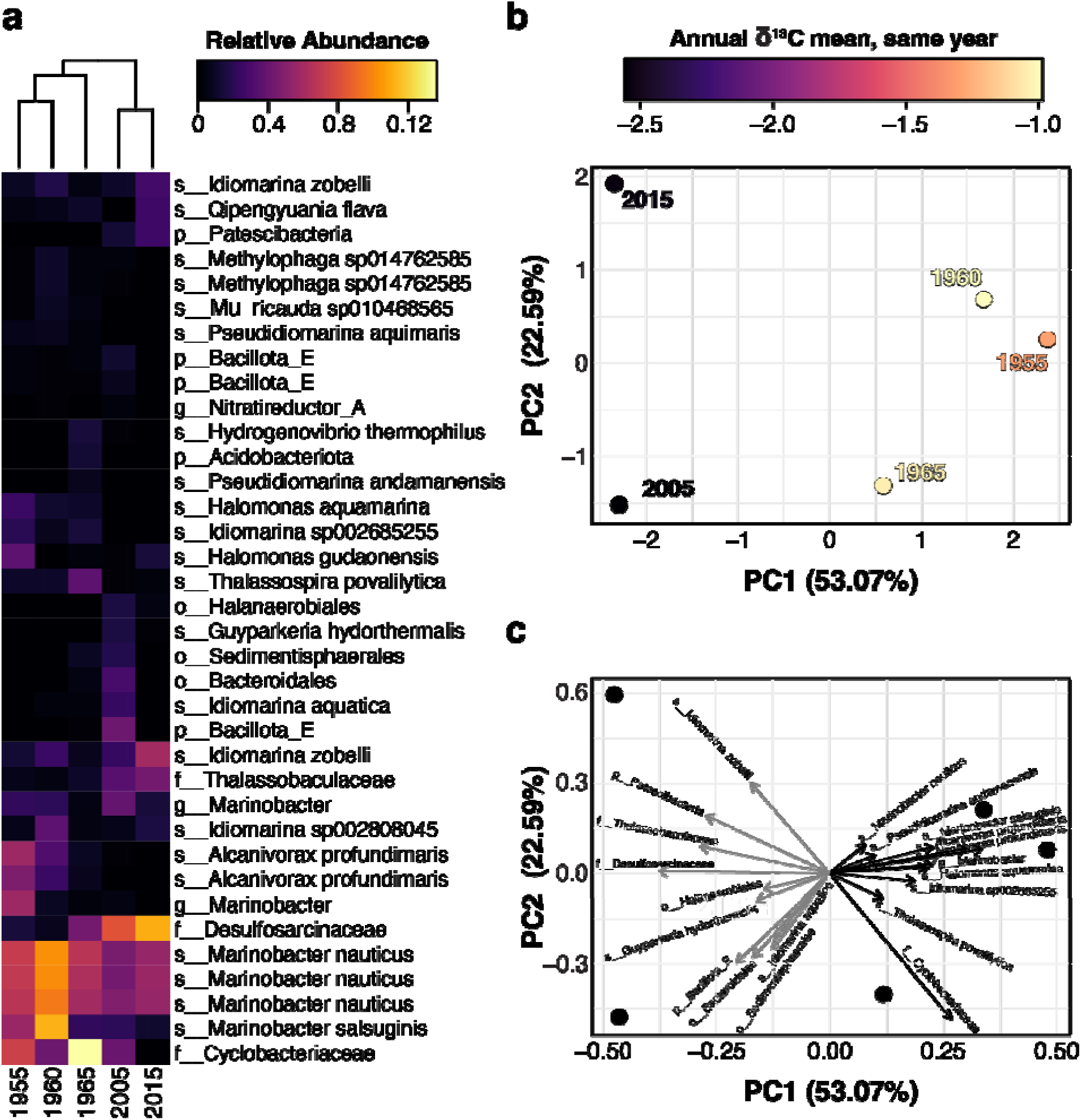
**(a)** Heatmap showing the relative abundance of MAGs recovered from the coral core according to the top right legend. Columns represent the core subsamples and rows the 36 MAGs. The finest taxonomic category assigned to each MAG by GTDB-Tk^151^ is shown on the right. Hierarchical clustering of columns (dendrogram on top) groups subsamples by chronology; the dendrogram clustering rows is not shown. **(b)** Principal Component Analysis (PCA) calculated from the relative abundance of the 36 MAGs. Each data point corresponds to a subsample (i.e., timepoint) and its color to the annual δ^13^C mean of the same year following the legend on top. **(c)** Loading plot of the PCA in **(b)** showing the MAGs driving the separation of subsamples along PC1 and PC2 bases on their contribution in both axes (Supplementary Table 6).

Functional analyses consisted of the identification of functional groups and profiling of the abundance of microbial metabolic pathways. Out of the 36 MAGs, 10 could be annotated as belonging to one or more functional groups using the Functional Annotation of Prokaryotic Taxa (FAPROTAX v1.2.9) database and software^59^ (Figure 4a; Supplementary Table 5). From these, hydrocarbon degradation is the only function that appears to have decreased over time with its highest abundance in 1955, whereas methanol oxidation and methylotrophy peaked in 1960. Several functions spiked in 1965, including chemoheterotrophy, dark sulfur oxidation, fermentation, hydrogen oxidation (i.e., knallgas bacteria), and ureolysis. Dark hydrogen oxidation and respiration of sulfur compounds were the only functions that increased over time. Microbial metabolic pathways were profiled using HUMAnN v3.0^60^. PCA separated the 1955 subsample from the rest (Figure 4b), driven by metabolic pathways involved in the production thiamine precursors (thiamine diphosphate salvage, thiamine phosphate formation from pyrithiamine and oxythiamine), peptidoglycan maturation, chlorophyll breakdown (phytol degradation), and amino acid biosynthesis (Supplementary Figure 9 and Supplementary Table 8). PC2 separated earlier from more recent timepoints also with an apparent connection to temperature (Figure 4b), driven by pathways related to the biosynthesis of polyamines (e.g., norspermidine), amino acids (L-arginine, L-tryptophan, L-isoleucine, L-alanine), fatty acids (5Z-dodecenoate) and cofactors (coenzyme A, flavin and heme B), and to metabolic energetics (homolactic fermentation, Entner-Doudoroff pathway, TCA cycle and glycogen degradation; Figure 4c and Supplementary Table 9).

**Figure 4.**
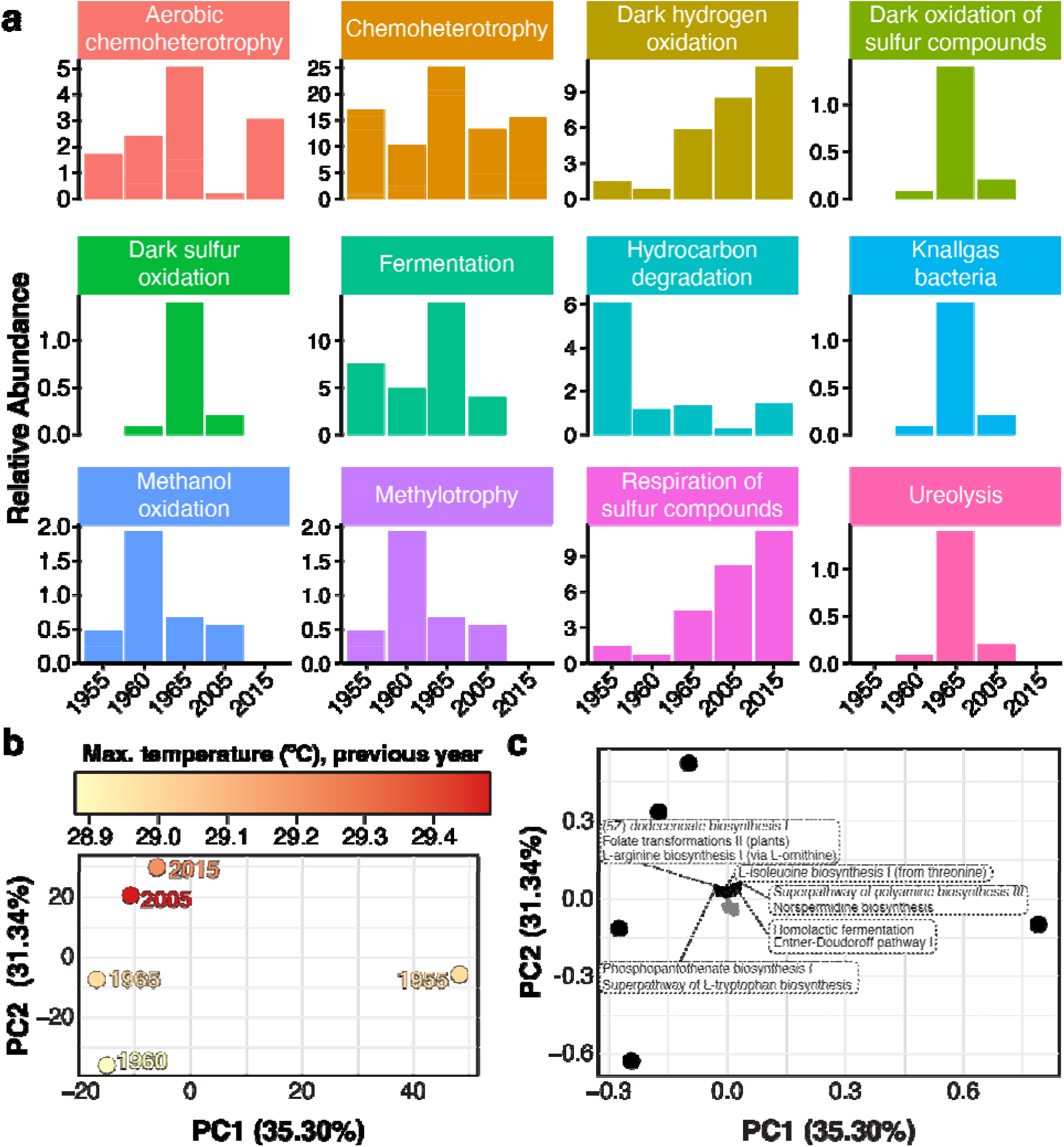
**(a)** Relative abundance of the FAPROTAX functional groups across timepoints. **(b)** Principal Component Analysis (PCA) calculated from the HUMAnN3 pathway abundances (see *Experimental procedures*). Each data point corresponds to a subsample (i.e., timepoint) and its color to the maximum annual temperature of the previous year following the legend on top. **(c)** Loading plot of the PCA in **(b)** showing the top 10 pathways contributing to PC2 in both directions (positive and negative). Labels for those contributing to the PC2 positive direction are shown, for detail on those contributing to the negative direction see Supplementary Table 9.

### Higher abundance of viral sequences is consistent with increase in predicted virulence

A total of 945 sequences (adding up to 6,332,358 bp) from metagenomic assemblies were predicted to be of viral origin with either high or medium confidence (see *Experimental procedures*; Supplementary Table 10). PCA separated 1965 and 2015 from the other timepoints (Figure 5a), most likely due to the high abundance of *Privateervirus* and Suoliviridae in the former, and of Caudoviricetes in the latter (Figure 5b, Supplementary Figure 9). Though no apparent links with temperature were observed (Supplementary Figure 10), the separation of 2015 might be associated with a high variation in water flow from Canal del Dique, as indicated from the δ^13^C proxy (Figure 5a). We were only able to explore fluctuation between predicted virulent (i.e., lytic) and temperate (i.e., lysogenic) lifestyles for the Order Caudoviricetes (Figure 5c), the only taxon present across all five timepoints. The virulent lifestyle peaked in 1965, which coincides with the highest abundance of that order across timepoints.

**Figure 5.**
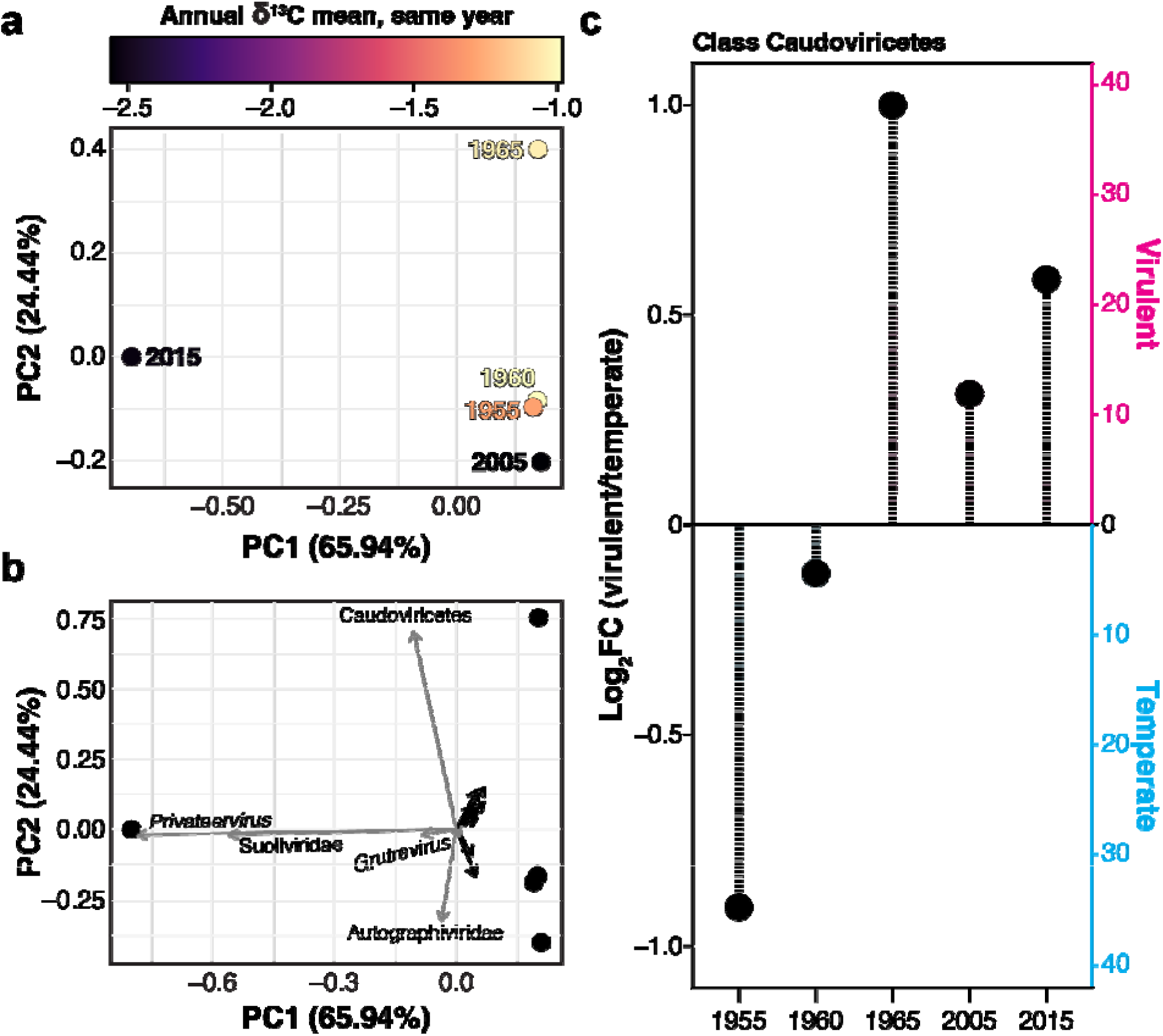
**(a)** Principal Component Analysis (PCA) calculated from the relative abundances of viruses predicted using PhaBox (see *Experimental procedures*). Each data point corresponds to a subsample (i.e., timepoint) and its color to the annual δ^13^C range (proxy of water flow from Canal del Dique) of the same year following the legend on top. **(b)** Loading plot of the PCA in **(a)** showing the top 10 pathways contributing to PC1 in both directions (positive and negative). Labels for some of the viral taxa with the largest weight on PC1 are shown (Supplementary Table 8). **(c**) Lifestyle prediction, either virulent (i.e., lytic) or temperate (i.e., lysogenic), for viral sequences classified within order Caudoviricetes across timepoints. The bar plot shows the number of sequences for each lifestyle (right *y*-axis). The lollipop plot shows the log_2_-fold change of the virulent to temperate ratio of the number of sequences (left *y*-axis).

## Discussion

In this study, *cora*DNA was successfully recovered from a skeletal core of the Caribbean coral *Orbicella faveolata*. Though sequence data were limited for several of our timepoints (Figure 3b), evidence of typical aDNA damage (Supplementary Figure 5) and undetectable contamination from environmental sources (Supplementary Figure 8) support the reliability of findings derived from timepoints with sufficient data. Endogenous DNA was detected from various members of the coral holobiont, including the animal host, the dinoflagellate photosymbionts, endolithic algae and associated microbiome (Figures 3–5). Coral skeletal and geochemical data enabled the assignment of ages spanning six decades. We used historical environmental records and geochemical data from the same core to explore links between past environmental conditions and changes in the coral holobiont, which we discuss below.

### Algal symbiont shuffling may be associated with Canal del Dique discharge

Varadero Reef is a prime example of a long-term impacted reef because of local human activities. Corals in this reef have been exposed to high-sediment loads since 1954 to the present due to perturbations caused by freshwater discharges from the Magdalena River via the human-made Canal del Dique distributary channel (144 × 10^6^ solids into Cartagena Bay each year) on a far greater scale than other Caribbean reefs^53,54^. Historical records from the Cartagena Bay in Colombia showed a marked increase of surface seawater temperature annual maxima and minima since the 1950s (Figure 2e and Supplementary Table 4). Concurrently, the skeletal δ^13^C suggests an increase, and higher annual variation, in the water flow from Canal del Dique in the more recent timepoints. This observation coincides with increased water discharge and sediment load for Canal del Dique in the 2000s^61^. Potential impacts of both temperature and water flow in the *O. faveolata* holobiont are thus compounded.

Both rise of temperature and water flow could have caused algal symbiont shuffling. Though *Orbicella faveolata* is known to host members of four distinct Symbiodiniaceae genera (*Symbiodinium*, *Breviolum*, *Cladocopium*, and *Durusdinium*)^62^, often with co-occurrence of different ones in the same colony^63^, the detection of some genera at certain timepoints is likely spurious. For example, the higher diversity and evenness of Symbiodiniaceae in 1955 (Figure 2d) could be explained by substitutions resulting from DNA damage (Supplementary Figure 2), which might introduce substitutions in the sequence reads that cause them to align to the wrong genome reference. The available genome datasets used here as references might be another source of artifacts, in that reads would align to the most similar sequence and not to the one they originated from. Reference biases and spurious mappings are common issues in ancient and historical DNA datasets^64^ that should be kept in mind. Another intriguing example is the presence of *Durusdinium*. This genus is endemic to the Indo-Pacific^20^ and its introduction in the Caribbean is thought to have been driven by a mass coral bleaching event in 2005^65^, so its detection in earlier timepoints is likely spurious. Nevertheless, its presence in 2015 is consistent with its quick spread through the Caribbean due to its thermotolerance and the recurrent heatwaves since its introduction^66^. Members of *Fugacium* are typically non-symbiotic or symbionts of foraminifera^20^, and there are no records of this genus being hosted by *Orbicella* corals. Even though *O. faveolata* can host *Breviolum*, its relative abundance here was two orders of magnitude lower than that of the other genera (Figure 2d,e). We thus focus on the most prevalent genera across timepoints, i.e., *Symbiodinium* and *Cladocopium*. Typically, *Symbiodinium* is found in either shallower *O. faveolata* colonies, or in parts of the colonies with the most exposure to light. Conversely, *Cladocopium* symbionts in *O. faveolata* occur at bigger depth or parts of the colonies with low light exposure^67^. We thus hypothesize that the observed fluctuations in relative abundances fluctuation depicts symbiont shuffling from a *Symbiodinium*- to a *Cladocopium*-dominated colony, driven mainly by water discharge from Canal del Dique. Indeed, sediment discharge from Canal del Dique has changed the optical properties of the seawater at Varadero Reef to resemble a reef at a greater depth^25^. In addition, *Cladocopium* symbionts associated with *Orbicella* spp. corals have previously been associated with high levels of sedimentation^63^.

### Microbial shifts linked to pollution and bleaching

The microbial taxa recovered through *cora*DNA (Supplementary Table 5) consisted mostly of common coral endolithic bacteria^68–70^. The *O. faveolata* microbiome was dominated by MAGs belonging to *Marinobacter* in early timepoints, 1955 and 1960 (Figure 3a). Members of this genus commonly establish mutualisms with dinoflagellates and other microalgae by mediating iron intake^71^. They are also part of the core microbiome of Symbiodiniaceae^72^, supporting their growth through metabolic exchange^73^. Members of this genus are known for their capacity to breakdown volatile hydrocarbons and their use for bioremediation of oil spills has been proposed^74,75^. However, the abundance of functional groups with the capacity to degrade hydrocarbons in 1955 (Figure 4a) is not related to the abundance of *Marinobacter* (Figure 3a), since none of the *Marinobacter* MAGs were classified within any functional group (Supplementary Table 5). The independent observations of *Marinobacter* and of the hydrocarbon degradation functional group point towards hydrocarbon pollution at Varadero Reef in the 1950s. Indeed, Cartagena Bay experienced substantial industrial development during this time, with an oil refinery starting operations, often with discharge of untreated organic matter, in 1957^76^.

Cyclobacteriaceae was the dominant taxon in 1965 (Figure 3a). These anaerobic bacteria are commonly found in corals^77^, where they are part of the endolithic community and potentially symbionts of *Ostreobium*^78^. Cyclobacteriaceae have been proposed as a bioindicator of high toxicity and heavy metal pollution^79^. The spike of this family in 1965 coincides with high mercury concentration in Cartagena Bay as result of the operations of a chlor-alkali plant^80^. The increased availability of environmental mercury can promote metabolic synergy between sulfur-reducing and -oxidizing bacteria, which might explain the increased abundance of dark-sulfur oxidation at this timepoint as well^81^ (Figure 4a). Such a bacterial interaction enhances mercury methylation, a process in which Cyclobacteriaceae might play a role^82,83^.

Desulfosarcinaceae is a family of anaerobic sulfate- and nitrate-reducing bacteria that proliferate in high-nutrient environments^84,85^. Its dominance in recent timepoints (2005 and 2015; Figure 3a) partly caused the increased abundance of functional groups able to perform dark hydrogen oxidation and respiration of sulfur compounds (Figure 3a and Supplementary Table 5). Desulfosarcinaceae may affect larval settlement and performance of *O. faveolata*^86^, but they can also contribute to further the microbial dysbiosis in bleached corals^87^. Its dominance in recent timepoints (Figure 3a), correlated with increased seawater temperature (Figure 3b), might be thus a sign of distress related to the 2005^88^, 2010^89^ and 2014-2015^90^ mass bleaching events in the Caribbean. Alternatively, the abundance of Desulfosarcinaeae could have been caused by enrichment of organic matter resulting from Canal del Dique sedimentation^80,91^.

### Proliferation of phages through virulence triggered by environmental stress

The viral sequences predicted from metagenome assemblies encompassed a great diversity of taxa (Supplementary Figure 10), including some that are commonly found in coral holobionts: families Ascoviridae, Mimiviridae and Phycodnaviridae (class Megaviricetes); family Baculoviridae (class Naldaviricetes); family Poxviridae (class Pokkesviricetes); and family Retroviridae (class Revtraviricetes)^92–94^. Nevertheless, class Caudoviricetes was the only taxon consistently found across the five timepoints, and the one explaining their differences. For example, 1965 distinguished itself from other timepoints because of the high abundance of phages in this class. The increase of Caudoviricetes could be explained by either the proliferation of their hosts or by replication of the viral particles. The dominant bacterial taxon in 1965 was family Cyclobacteriaceae (Figure 3a), but the most abundantly predicted phages hosts belong to Enterobacteriaceae (Supplementary Figure 12), a family not recovered by our MAGs (Supplementary Table 5). The increase in abundance of Caudoviricetes in 1965 coincides with the highest ratio of virulent relative to temperate sequences (Figure 5c), making viral replication through a lytic switch, in response to environmental stress, a more likely scenario. Caudoviricetes also have been reported in higher abundance in bleached corals^95^. The abundance of *Privateervirus* and Suloiviridae (both members of Caudoviricetes) might be another signal of distress, similar to Desulfosarcinaceae (see above).

### Technical limitations

Some technical restrictions of our novel approach warrant conservative interpretation of the results and represent room for improvement. Future studies combining *cora*DNA and metagenomic approaches would benefit from more intense and thorough sampling. This includes sampling the appropriate controls for decontamination^96^. In this case, appropriate controls would include samples from the surrounding environment (collected simultaneously with the coral cores) and from the extant live tissue of the target colony. A more intense sampling should also comprise biological replicates, which would enable statistical testing and strengthen conclusions^97^. Historical and ancient DNA have inherent sampling complications, such as limited access to biological material and frequent fall-through of samples due to little to no DNA recovery^98^, such as in this study. Another limitation specific to *cora*DNA is the availability of massive or bouldering colonies with continuous, regular growth along a uniform axis^99^. Therefore, a strategy in which corals are sampled to be simultaneously processed for geochemical and *cora*DNA analyses will enable tighter links to be drawn between environmental and holobiont changes. A final consideration is to budget for deeper sequencing. Deep sequencing can improve analysis of samples with low DNA yield and help recover low-abundance taxa (e.g., *Ostreobium* in this study) more consistently, and thus to identify contaminant microbes, which in turn provides more confidence in the representation of the actual microbiome^100^.

### Integrating coraDNA, genomics and geochemistry: a new field

This study is the proof-of-concept of the power of integrating sclerochronology, *cora*DNA, metagenomics and geochemistry techniques to reconstruct the response of coral holobionts to past environmental changes. This multi-pronged study lays the foundation for investigations to provide deeper and more-comprehensive insights of coral holobionts and reef communities over time (Figure 6). Here, surveyed holobiont members were restricted to the coral host, dinoflagellate symbionts, and associated bacteria and viruses. Nevertheless, this approach could be used the explore temporal and environmental dynamics of other more-elusive members of the holobiont in the future, such as endolithic algae^101^, protists^102^, fungi^103^, and archaea^104^. Reconstructing shifts in holobiont microbial communities across space and time over the lifespan of a coral colony will enable us to track taxa that have survived and prevailed over major environmental insults. With the increasing interest in probiotic treatments to help corals cope with climate change^105^, *cora*DNA can help guide targeted microbial culturing for this purpose in our quest to help reefs survive.

**Figure 6.**
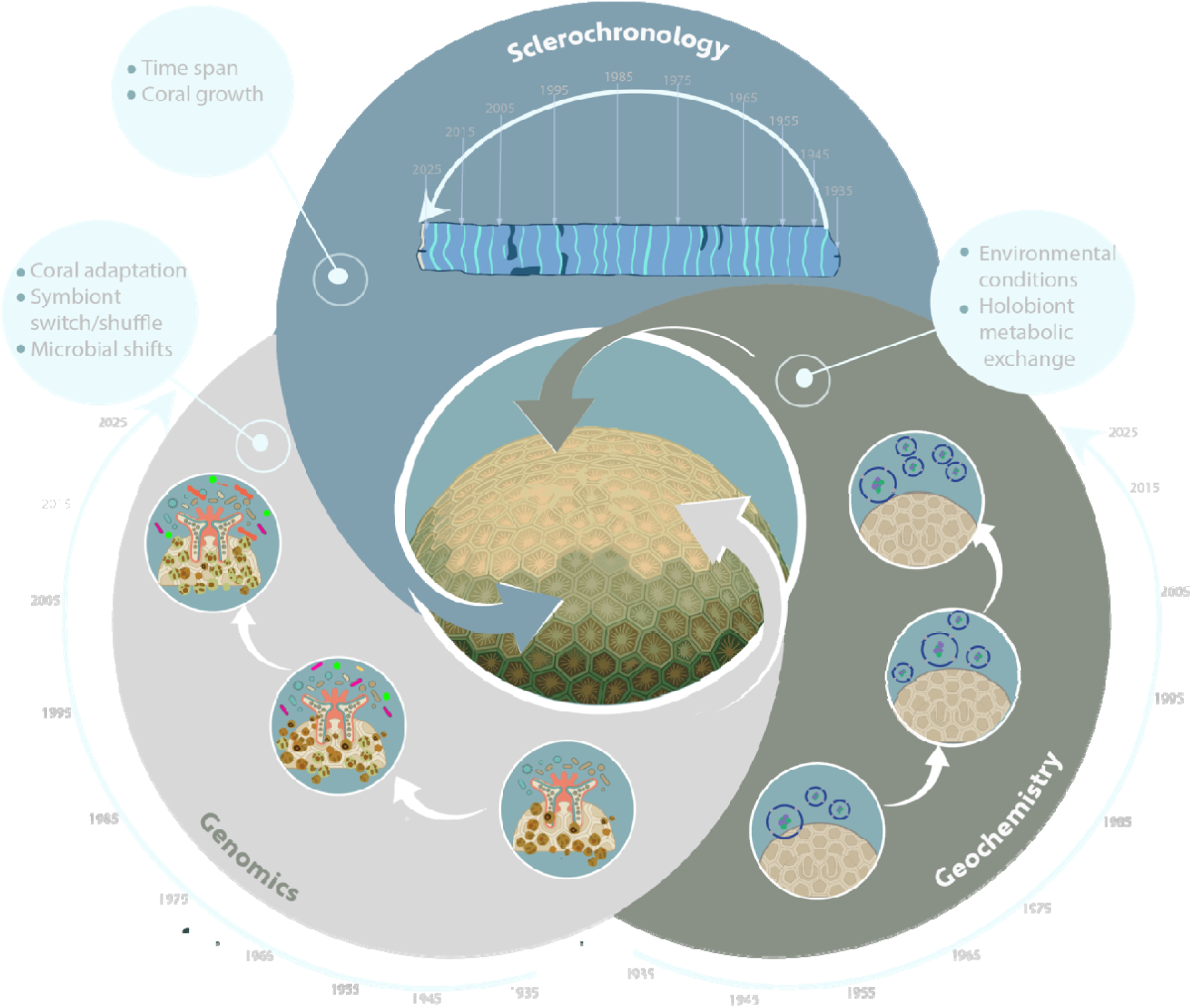
Multi-pronged approach–incorporating sclerochronology, genomics and geochemistry–to assess coral holobiont dynamics over time and to reconstruct historical responses to environmental change.

Likewise, additional techniques can provide further insights into interactions of the coral holobiont with its past environment. Some omics can uncover rapid response mechanisms to environmental stress and fluctuations. Reconstruction of DNA methylation maps in historical and ancient samples can reveal changes and in gene regulation over time^106^. Plasmidomics (i.e., the characterization of the overall plasmid content in a certain environment^107^) can identify horizontal gene transfer playing essential roles in microbial evolution^108^.

Investigations targeting specific taxa, genes, or functional traits are also in the horizon. Whole-genome sequencing of specific taxa can now be achieved through enrichment approaches that usually involve the design of DNA baits^109,110^. This could allow, for instance, tracking the evolution of members of the coral microbiome with greater detail over time. Analogously, one could target a specific gene or gene set to study gene evolution. Likewise, the temporal progression of specific functional features could be interrogated, such as the production of a given metabolite or a pathway involved in nutrient cycling.

Beyond the coral holobiont, other technologies could be applied to explore the temporal succession of entire reef ecosystems. Sedimentary ancient DNA (sedaDNA) of reef substrates in combination with metabarcoding–comprising markers for fish, invertebrates, microalgae, and other organisms–could provide an overall picture of diversity and transitions of reef communities over time^111^. Like in this study, metagenomics and paleoclimatology techniques could also be used to reconstruct past microbial shifts and environmental fluctuations^111^.

This is only a brief overview of all the potential that our approach holds to explore coral holobionts and reef ecosystems from a whole new perspective. It is our hope that the knowledge generated through this kind of research will help predict responses of coral holobionts to the ongoing global environmental change, and in turn inform the conservation of coral reef ecosystems.

## Experimental procedures

### Coral sampling

A colony of *Orbicella faveolata* was sampled from Varadero Reef, southwest of Cartagena Bay, ∼6 km West to the Canal del Dique outlet (10° 18′ 23.3″ N, 75° 35′ 08.0″ W, Figure 1a-c). A core of 6 cm diameter was collected along the growing axis of the skeleton at 3–5 m depth by SCUBA, following a previously described method^112^ with a modified version of the coring device that consists of an underwater pneumatic hand drill supplied with compressed air coupled to customized stainless-steel pipes with a coring-head with tungsten teeth (Figure 1e). The affected live coral tissue (25-30 cm^2^) was covered with a polystyrene ball and epoxy to avoid intrusion of bioeroder organisms. The core was rinsed with fresh water several times to remove sediment particles and then oven-dried for 24–48 h at 50–60 °C.

### Environmental data

Monthly water flow records (1981‒2015) from Canal del Dique at the Santa Helena station (ID 29037450) were obtained from the *Instituto de Hidrología, Metereología y Estudios Ambientales* (IDEAM; ideam.gov.co). Sea surface temperature and precipitation (9–11°N, 283–285°E; 1951–2020) records retrieved from the HadISST1^113^ and CRU TS v4.08 (crudata.uea.ac.uk/cru/data/hrg/cru_ts_4.08) datasets, respectively, and processed and downloaded using Climate Explorer (climexp.knmi.nl/start.cgi). Annual metrics were derived from the monthly records, including maximum and minimum values, mean, and range (i.e., maximum – minimum). Both monthly records and annual metrics were tested for Pearson correlation against geochemical measurements using R v4.5.2.

### Geochemistry and age model

Processing of the core for elemental, isotopic, and sclerochronology analyses was done in the PACE Laboratory, at the University of Michigan’s Earth and Environmental Science Department. Slabs (5 mm thick) were taken from the center of each section of the coral core. Samples were micro-milled at 1-mm resolution along the axis of maximum growth from a transect 4 mm wide and 2 mm deep. These homogenously powdered samples were divided and measured for elements (Sr/Ca, Mg/Ca, Ba/Ca) and stable isotopes (δ^18^O, δ^13^C), following established protocols^43^. Briefly, for elemental values, a sample of 0.4–0.6 mg was measured using a Thermo iCAP 7400 inductively coupled plasma optical emission spectrometer (ICP-OES). Isotopic samples of 40–60 μg were analyzed using a Thermo Delta V Plus mass spectrometer, coupled to a Kiel IV carbonate preparation system.

Coral luminescence was estimated by digitally extracting the filtered intensity of red, green, and blue colors from luminescent images along the same transects used for geochemistry using Fiji^114^. We then calculated the green-blue ratio and interpolated the series to have the same sample resolution as the geochemical record. Coral density was estimated using 3D X-ray scanning of the coral slabs using a micro-CT scanner (Nikon X-Tek XTH225ST). Density was estimated from Hounsfield units (HU), which were extracted from a 2-mm thick slab average along the same transects used for geochemistry using Fiji.

An age model was constructed for the coral core using annual cycles in the quasi-sinusoidal Sr/Ca, Ba/Ca, δ^18^O, and luminescence signals. The coldest month (February or March, based on the coolest HadSST for a given year) was assigned to the subsample with the most positive Sr/Ca, δ^18^O and the lowest Ba/Ca and luminescence. The warmest month (August, September, or October, based on warmest HadSST for a given year) was assigned to the subsample with the lowest Sr/Ca, δ^18^O and the highest Ba/Ca and luminescence. Using these tie points, we assigned ages to each sample and then interpolated the geochemical, luminescence, and density records to monthly resolution.

### DNA extraction and next-generation sequencing

Annual growth bands in *O. faveolata* are about 1 cm thick, subsamples (∼1.5 cm^2^ squares) were taken in intervals of 5 cm (i.e., every ∼5 years), resulting in 13 subsamples (Supplementary Figure 1). The subsamples were furthered processed at the Penn State Ancient DNA Laboratory, where no previous work with corals had been undertaken before. Negative controls were kept throughout the sample processing. All surface areas and subsampling equipment were sterilized after each sample using a 10% bleach solution, rinsed with 70% ethanol, and followed by UV-irradiated distilled water. A Dremel with a diamond-cutting wheel was used, on a low-speed setting, to fraction each sample into a subsample of ∼50-105 mg, ensuring the retrieval of the original square’s inner core and avoiding the outer sides “exposed” during transport. The 13 subsamples were then mechanically pulverized and transferred to 2 ml tubes for subsequent DNA extraction.

DNA extraction was based on a customized protocol for ancient samples^115–117^, proven successful in studies focused on organisms that precipitate calcium carbonate, like corals^50^ and mollusks^117,118^. For initial demineralization, 1 ml of 0.5 M EDTA (pH = 8) was added to each subsample, incubated at 22 °C (room temperature) and shaken at 1,000 rpm in a thermomixer for 24 h. The tubes were then centrifuged at 12,800 *g*, and the supernatant transferred to a new 2 ml tube and stored at -20 °C. For enzymatic digestion, 1 ml of extraction buffer (0.5 M pH 8 EDTA, 0.5% sodium dodecyl sulfate (SDS), 0.25 mg/ml proteinase K) was added to each pellet, followed by vortex resuspension. The samples were then incubated at 37 °C and shaken at 1,000 rpm in a thermomixer for 24 h. After a 12,800 *g* spin, the supernatant was pooled with the former one. The combined DNA supernatant (2 ml) was poured into 50 ml Falcon tubes with 5 volumes of QIAquick PB buffer, 400 µl 3 M sodium acetate (pH 5.2) and 5 µl Pellet Paint. The extraction solution was added to a QIAquick spin column that was fitted with a Zymo-Spin V extender and then secured in a vacuum manifold. After filtering, the QIAquick spin columns were transferred to 2 ml collection tubes for a 3,330 *g*-dry spin, followed by a 750 µl QIAquick PE buffer wash and centrifugation at 3,300 *g* for 1 min. The collected solution was discarded and a subsequent dry spinnned at 12,800 *g* for 1 min. DNA was eluted in two 25 µl portions of QIAquick EB at room temperature for 2 min prior to centrifugation at 12,800 *g* for 1 min. The purified DNA was then stored at -20°C. Two negative controls (extraction blanks) were included alongside the 13 subsamples.

Aliquots of the DNA extract (20 µl), the two extraction controls and a water blank were converted into double-stranded libraries suitable for Illumina platform sequencing following a modified protocol^39,40^. Blunt-end repair was performed without the removal of cytosine residues, followed by purification using the MinElute PCR Purification Kit (Qiagen) and heat deactivation at 80 °C for 20 min was done after adapter fill-in. The final libraries (40 µl) were used in the subsequent indexing amplification based a previous protocol^119^, using the KAPA HiFi HotStart ReadyMix (25 µl per reaction), the p7 primer (300 nM) incorporating a unique index for each library (5′-CAAGCAGAAGACGGCATACGAGATxxxxxxGTGACTGGAGTTCAGACGTGT-3′, where ‘xxxxxx’ is the unique barcode) and a universal IS4 index (5′-AATGATACGGCGACCACCGAGATCTACACTCTTTCCCTACACGACGCTCTT-3′). Libraries were amplified following the protocol: 2 min at 98 °C; 11 cycles of 20 s at 98 °C, 30 s at 60 °C and 30 s at 72 °C; final extension 5 min at 72 °C. Indexed PCR products were purified using SPRI beads (1 mg/ml Sera-Mag carboxylate-modified magnetic particles (GE Healthcare), 38% PEG-8000 (w/v), 1M NaCl, 0.05% Tween-20, 1 mM EDTA and 10 mM Tris-HCl, pH 8.0) with 70% ethanol washes and a final elution in 20 μl 0.5% Tween-Tris/EDTA buffer. DNA concentration for all samples was determined using a Qubit 3.0 dsDNA High Sensitivity Kit; libraries were pooled in equimolar ratios based on their corresponding concentration. Shotgun sequencing was done at the NYU Genome Technology Center on an Illumina NextSeq 500 platform (2 × 75 bp). All raw reads are available in SRA BioProject PRJNA1494103 (Supplementary Table 1).

### Assessment of coral holobiont dynamics over time

For assessment of coral holobiont composition over time, the demultiplexed sequence data were input to leeHom v.1.2.16^120^ to trim adaptors and merge paired-end reads with a minimum overlap of 11 bp into single-end reads using the “--ancientdna” option, which implements a Bayesian framework to reconstruct typical short aDNA fragments. Merged single-end reads were further trimmed and filtered using trimmomatic v0.39^121^, keeping reads with an average quality ≥30 and at least 30 bp long (Supplementary Table 1).

We then aligned all reads pooled subsequently to genome references (Supplementary Table 2) of *O. faveolata*^122^, Symbiodiniaceae^32,123–131^, *Ostreobium quekettii*^132^, and to the final MAGs (Supplementary Table 3; see below), using *bwa aln* v0.7.18^133^ with settings (*-l 1024 -n 0.01 -o 2*) optimized for aDNA data^134,135^ and only the unaligned reads from the previous step; the number of aligned reads was determined using SAMStat 2 v2.2.3^136^. Reads not aligning to any of the genome sequences were used as query against the pre-formatted BLAST nucleotide (nt) database v5 using BLASTn v2.14.1+^137^, allowing only one top hit and the highest scoring pair (*e*-value ≤ 1×10^−25^); taxonomy identity was visualized with a sunburst plot produced by ExTaxsI vX^138^. Read count tables for each MAG, coral host, and algal symbionts were generated using CoverM v0.7.0 (github.com/wwood/CoverM) for each timepoint and all pooled together.

Metagenomes from each of the five timepoints, assembled using the trimmed reads and MEGAHIT v1.2.9^139^, were screened for viral sequences using the PhaBOX2 web server^140^ (accessed in July 2025), which identifies putative viral contigs, classifies them based on taxonomy, and predicts potential hosts and lifestyle. Contigs predicted to be viral with “medium-” and “high-confidence” were used for subsequent analysis. The trimmed reads were aligned to the metagenome assemblies just as before (*bwa aln -l 1024 -n 0.01 -o 2*) and read counts for estimates of relative abundance were also generated using CoverM v0.7.0.

### DNA damage and metagenomic analyses

Raw, demultiplexed read sequence data from the core subsamples (Supplementary Table 1) were used to evaluate DNA damage. To assess fragment size distribution, read pairs from each subsample were aligned against the assembled metagenome-assembled genomes (MAGs; see below) using bwa v0.7.19^133^; fragment sizes were then calculated using SAMtools v1.22.1^141^. Signals of cytosine deamination were detected using subsamples with at least 5 M reads and PyDamage v0.72^142^. These subsamples were then input into the Earth Hologenome Initiative (EHI) bioinformatics pipeline v1.0 (github.com/earthhologenome/EHI_bioinformatics). Briefly, adapters were trimmed, and low-quality (PHRED ≥ 20) and short (<30 bp) reads were removed using fastp v0.23.1^143^. Sequences from eukaryotic members of the holobiont were removed by aligning the filtered reads to genome references of the coral host, and of Symbiodiniaceae and *Ostreobium* symbionts (Supplementary Table 2)^32,122–132^, using Bowtie2 v2.4.4^144^. The remaining “non-host” reads were used for assembly per timepoint separately and pooled together (i.e., six distinct assemblies) using megahit v1.2.9^139^. Raw reads were aligned to the contigs of each assembly using Bowtie2. The contigs of each assembly were binned using three programs: CONCOCT v1.0.0^145^, MetaBAT 2 v2.12.1^146^, and MaxBin 2 v2.2.6^147^. We used the *bin_refinement* tool from metaWRAP v1.3.2^148^ to obtain a set of optimized, non-redundant bins per assembly. We only retained bins with ≥70% completeness and ≤10% contamination from each assembly as estimated with CheckM v1.0.12^149^. The final metagenome-assembled genomes (MAGs) were obtained by dereplication of all bins from all assemblies using dRep v3.4.0^150^ at an average nucleotide identity (ANImf) threshold of 98%. Taxonomic assignment of the final MAGs was done using GTDB-Tk v2.3.2^151^ and GTDB release 214^152^.

### Assessment of contamination

We looked for potential microbial contaminants from environment sources by comparing the assembled coral MAGs (Supplementary Table 5) to available metagenomic data from the Varadero Reef and from the proximate Rosario Islands reef (Supplementary Table 6). Briefly, reads were quality-assessed with FastQC v0.11.9 (bioinformatics.babraham.ac.uk/projects/fastqc), and low-quality (PHRED ≥ 20) and short (< 50 bp) reads, as well as poly-G tails, were removed using fastp v0.23.4^143^; adapters were further removed, and reads further filtered (PHRED ≥ 30), using trimmomatic v0.39^121^. Metagenome assembly was conducted pooling samples of the same location and type using megahit v1.2.9^139^, and subsequently binned using MaxBin v2.2.7^147^; taxonomic assignment was done using GTDB-Tk v2.3.2^151^ and GTDB release 214^152^ (Supplementary Table 5).

### Functional analyses

Assignment of functional groups was done using FAPROTAX v1.2.10^153^, normalizing functional group read counts with total sum scaling (TSS). Metabolic pathway abundance was profiled using HUMAnN v3.9^60^ and the full ChocoPhlAn and UniRef databases (downloaded in January 2026) and subsequent analyses were done following the *Functional Profiling* tutorial of the SPAAM-Community^154^.

## Acknowledgements

R.A.G.P was supported by an Eberly Postdoctoral Research Fellowship, awarded by the Eberly College of Science at The Pennsylvania State University. This work was supported by the NSF Biodiversity on a Changing Planet program (grant 2449251 awarded to M.M., J.E.C., L. S.W. and G.H.P) and by an NSF Division of Integrative Organismal Systems core program (grant 2227070 awarded to M.M.). The core samples were retrieved and analyzed under the auspices of a collaborative research initiative between Pontificia Universidad Javeriana Cali and The University of Queensland led by M.L.V., A.M.M.G. and A.R.R. We thank Ana Saborio for her help designing Figure 6.

