## Supplementary Figures for "A paleogenomic and paleoclimatic approach to reconstruct historical responses of coral holobionts to anthropogenic change"

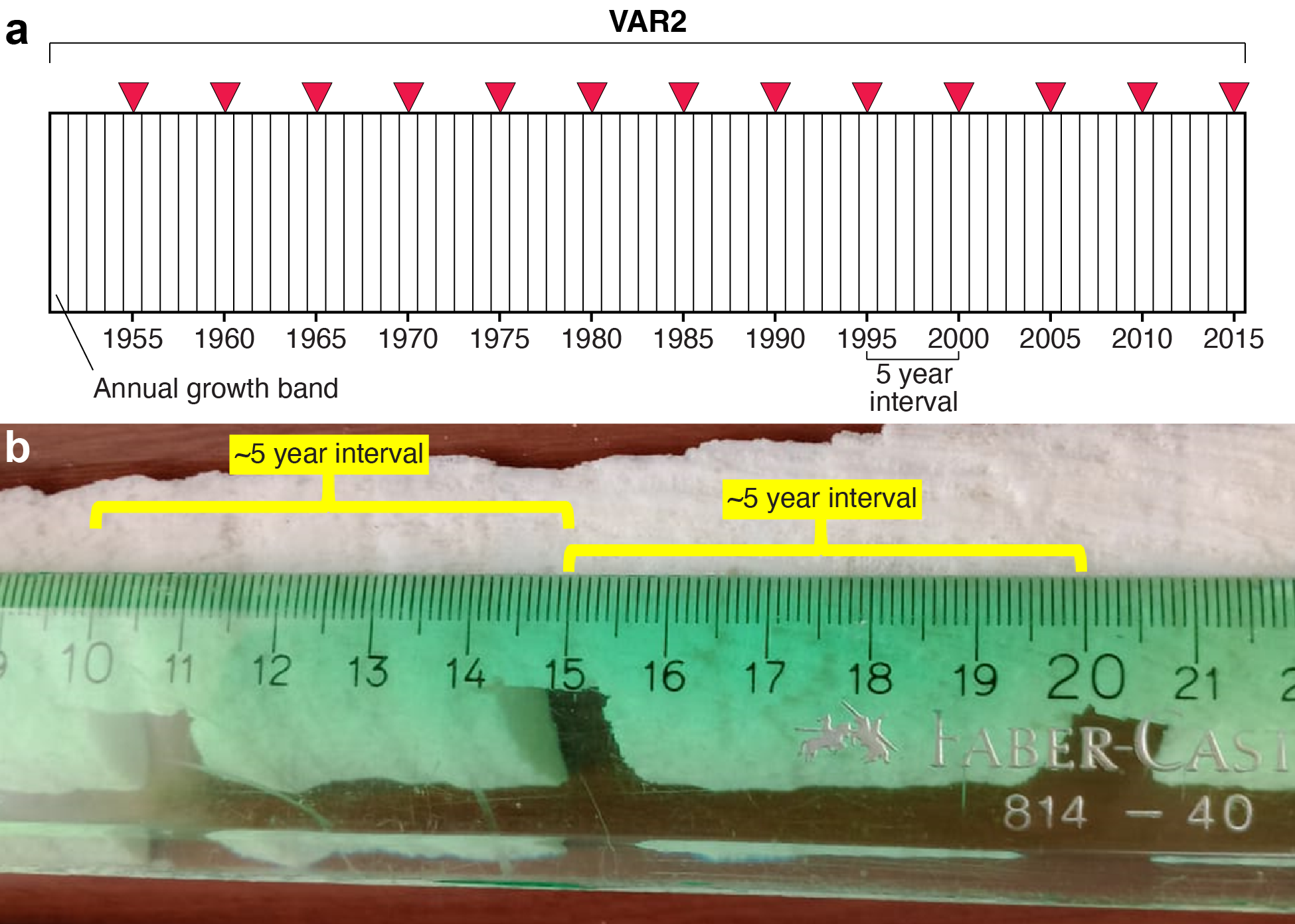

**Supplementary Figure 1.** **(a)** Diagram of the coral core subsampling strategy for *cora*DNA analysis. Red arrowheads indicate locations along the core where subsamples were taken. **(b)** Remains of the core after subsampling showing details about the ~5-year intervals (~5 cm) and subsampled squares along the bottom edge of the core (~1 cm^2^).

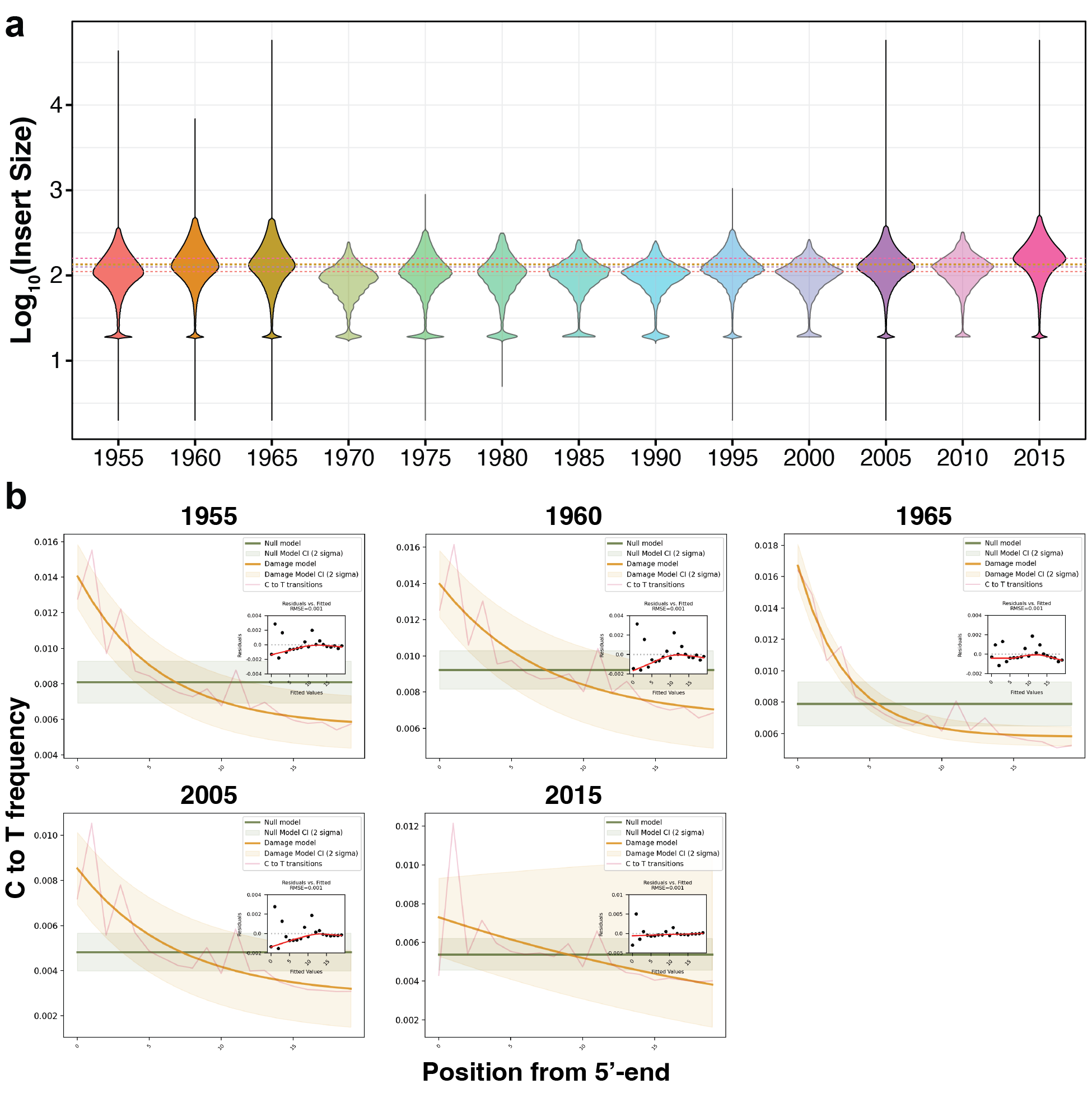

**Supplementary Figure 2. (a)** Fragment size distribution determined from processed read pairs from each subsample. Dotted lines correspond to the medians of subsamples with ≥5 M read pairs; the color of each line matches its violin plot. **(b)** PyDamage profiles of subsamples with ≥5 M read pairs.

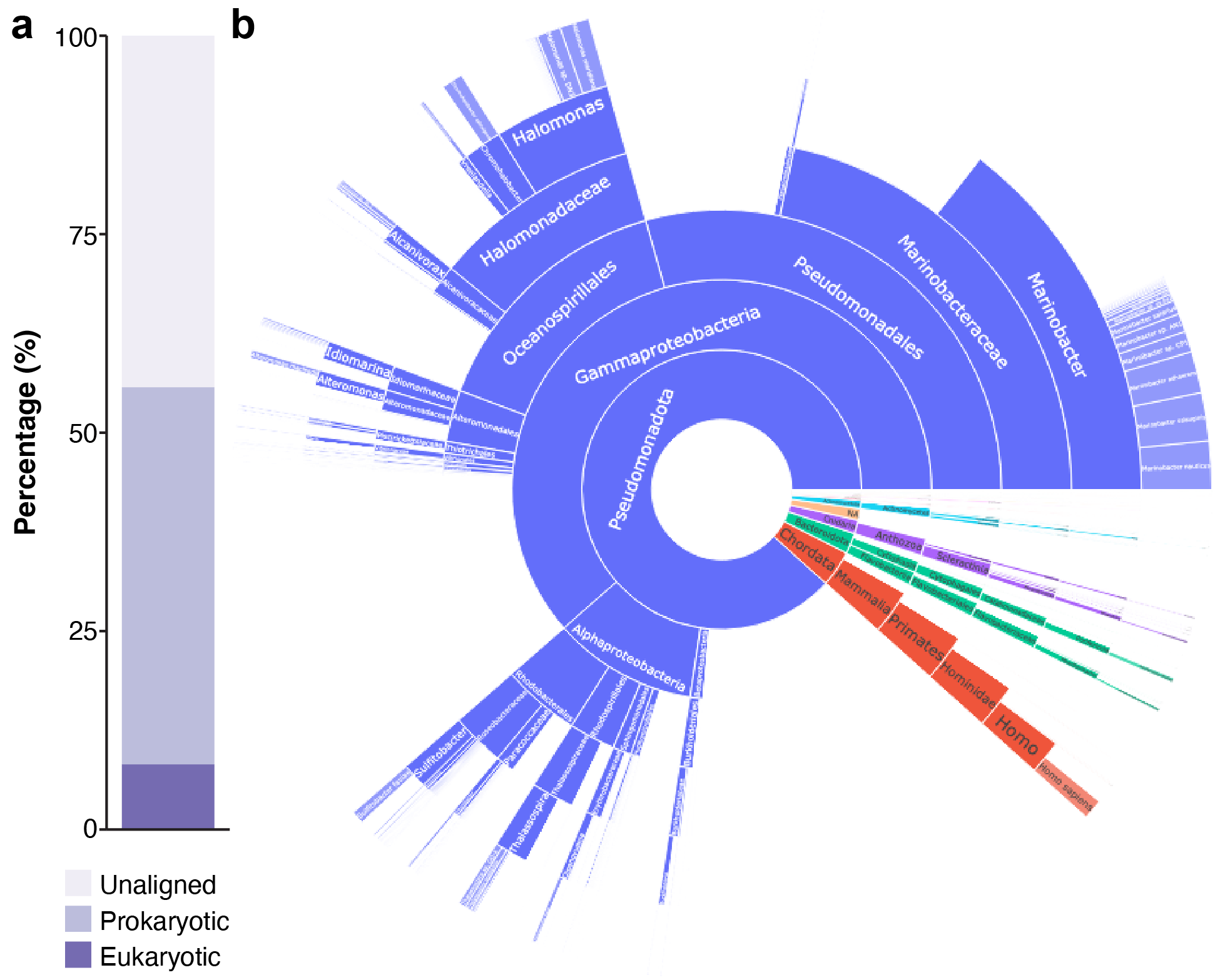

**Supplementary Figure 3. (a)** Breakdown of the total number processed of reads according to alignments against genome sequences of eukaryotic and prokaryotic members of the holobiont (see *Experimental Procedures*). **(b)** Sunburst plot with detail of the taxonomic identity of the unaligned reads in **(a)**.

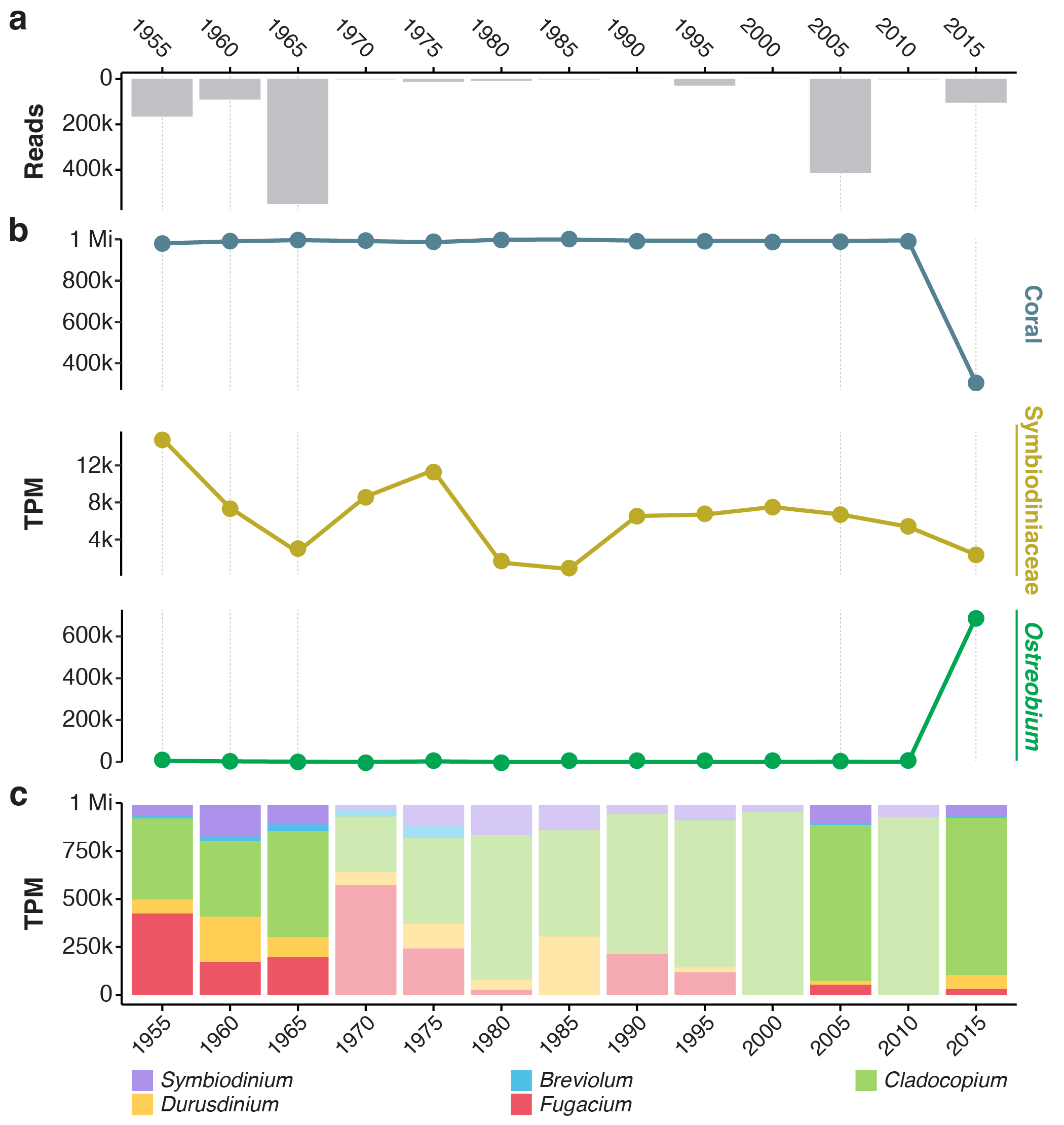

**Supplementary Figure 4. (a)** Number of processed reads that were used as input for alignments in **(b)** and **(c)**. **(b)** Relative abundance based on estimated Transcripts Per Million (TPM) of processed reads with high-quality alignments (MAPQ ≥ 30) to genome references (Supplementary Table 4) of the coral host (top), Symbiodiniaceae symbionts (middle) and *Ostreobium* (bottom) over time. **(c)** Same as in **(b)** but only alignments for Symbiodiniaceae are shown, broken down by genus. Subsamples with an initial sequencing depth ≥5 M reads are shown with a dashed gray line in the background in **(a)** and **(b)**; those with <5 M reads are represented as semi-transparent in **(c)**.

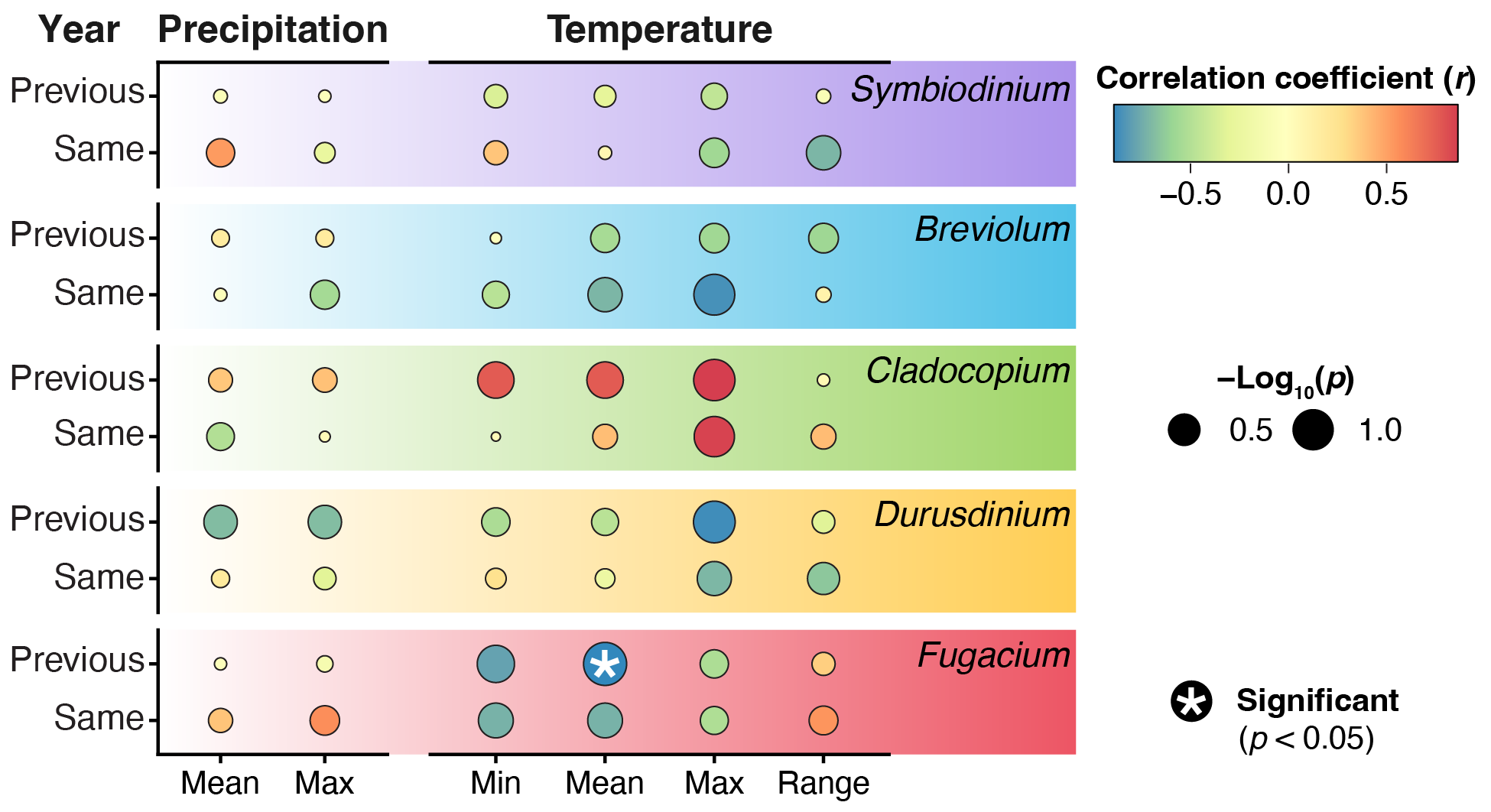

**Supplementary Figure 5.** Summary of Pearson correlation tests of relative abundances of Symbiodiniaceae genera with environmental variables (CRU TS precipitation and HadISST temperature records) in subsamples with enough read data. To assess potential effect of environmental variables on symbiont shifts, several yearly metrics were derived from monthly measurements (minimum, mean, maximum and the range of variation; *x*-axis) from the same year the relative of the symbionts was quantified, as well as from the year before (*y*-axis). The strength of the correlation is represented by the color of each data point and its statistical significance by the size following the legends to the right. Though most correlations were not statistically significant (*p* < 0.05), temperature exhibited the strongest ones. The abundance of 0 among the minimum yearly values of precipitation restricted its analysis to mean and maximum values. Water flow data from *Canal del Dique* were insufficient to perform the correlation tests.

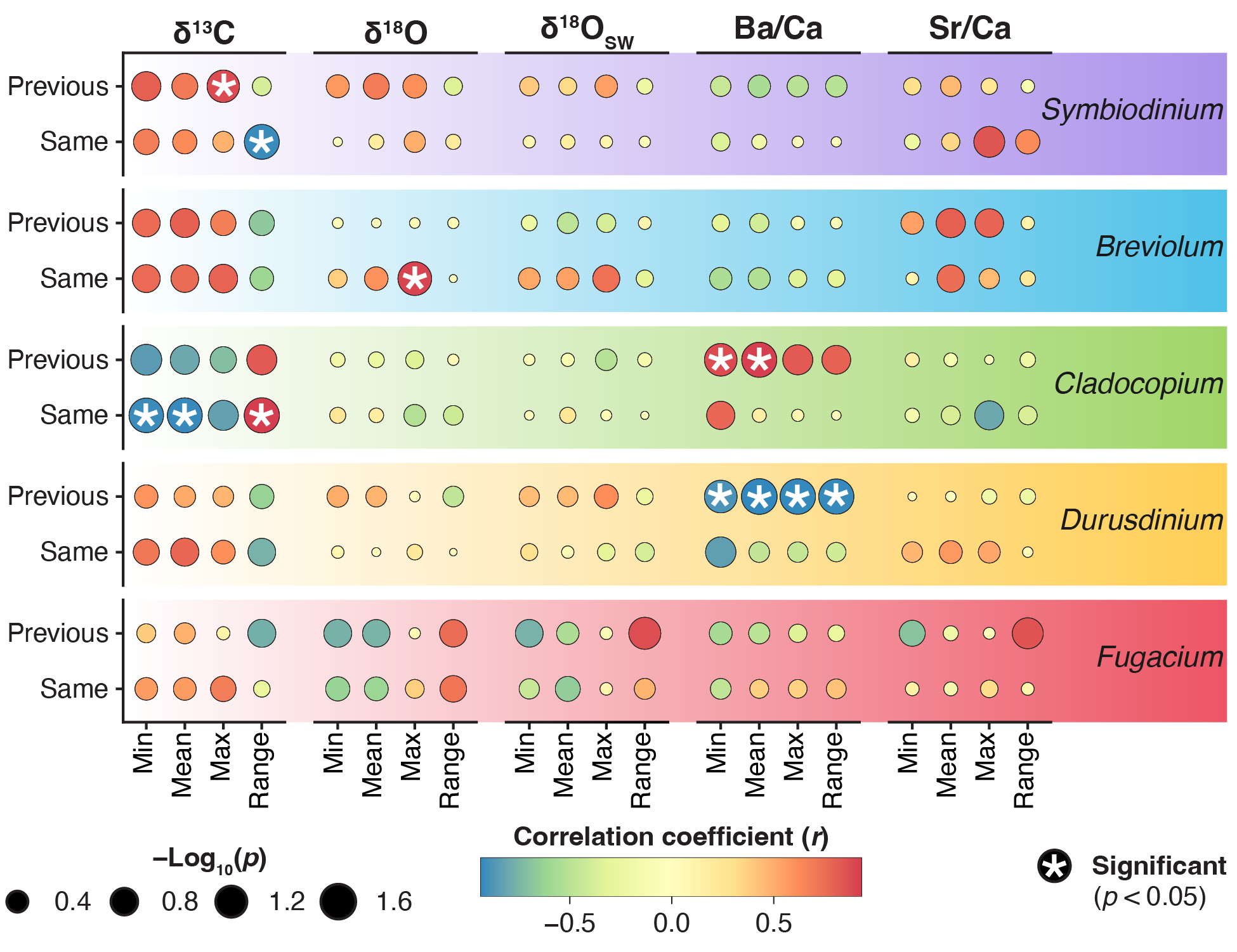

**Supplementary Figure 6.** Summary of Pearson correlation tests of relative abundances of Symbiodiniaceae genera with geochemical measurements in subsamples with enough read data. To assess potential effect of past geochemical measurements on symbiont shifts, several yearly metrics were derived from monthly measurements (minimum, mean, maximum and the range of variation; *x*-axis) from the same year the relative of the symbionts was quantified, as well as from the year before (*y*-axis). The strength of the correlation is represented by the color of each data point and its statistical significance by the size following the bottom legends.

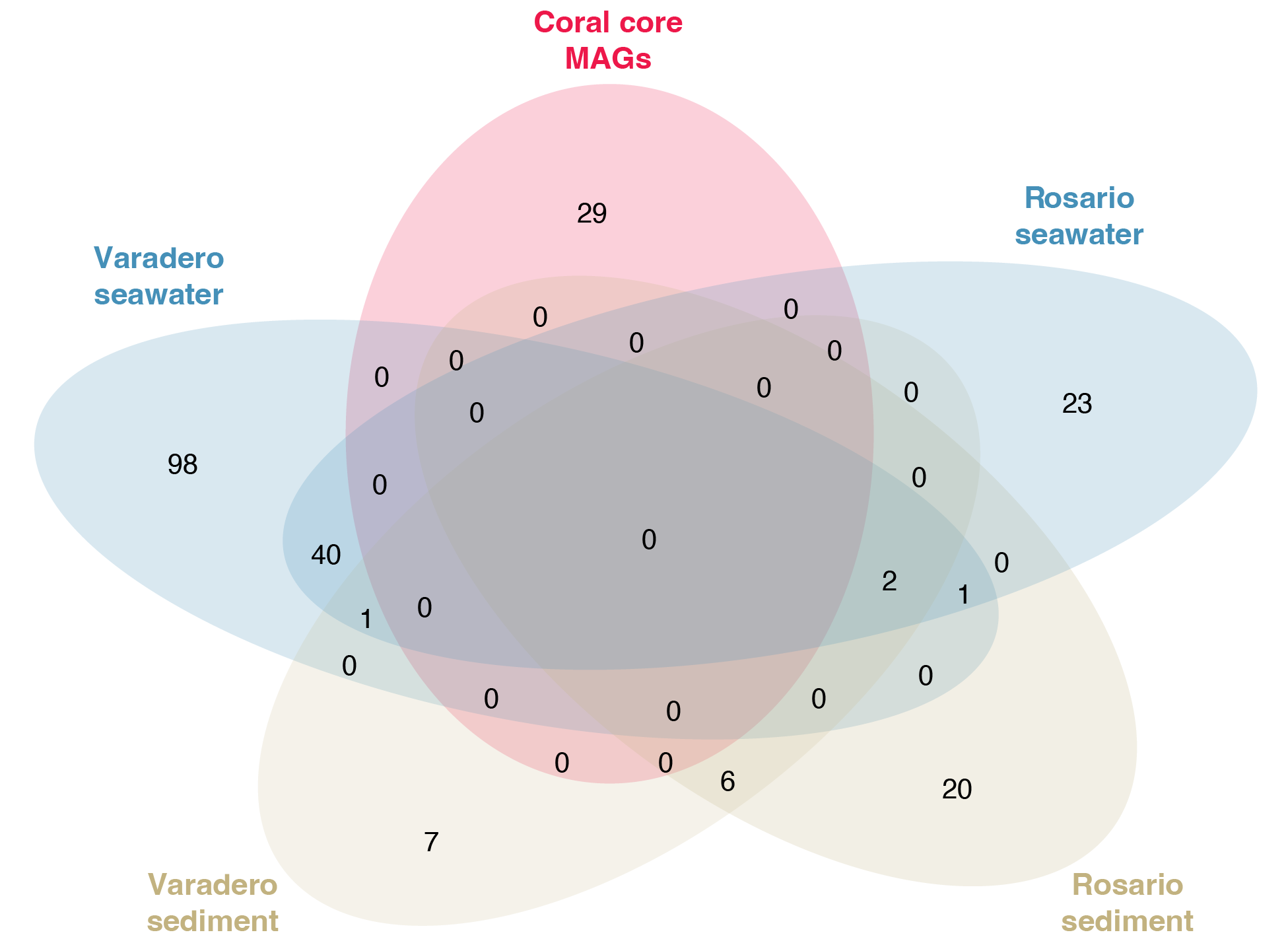

**Supplementary Figure 7.** Venn diagram showing the number of taxa overlapping between generated MAGs from the coral core and from the environment (seawater and sediment) at the sampling site (Varadero Reef) and a nearby reef (Rosario Islands).

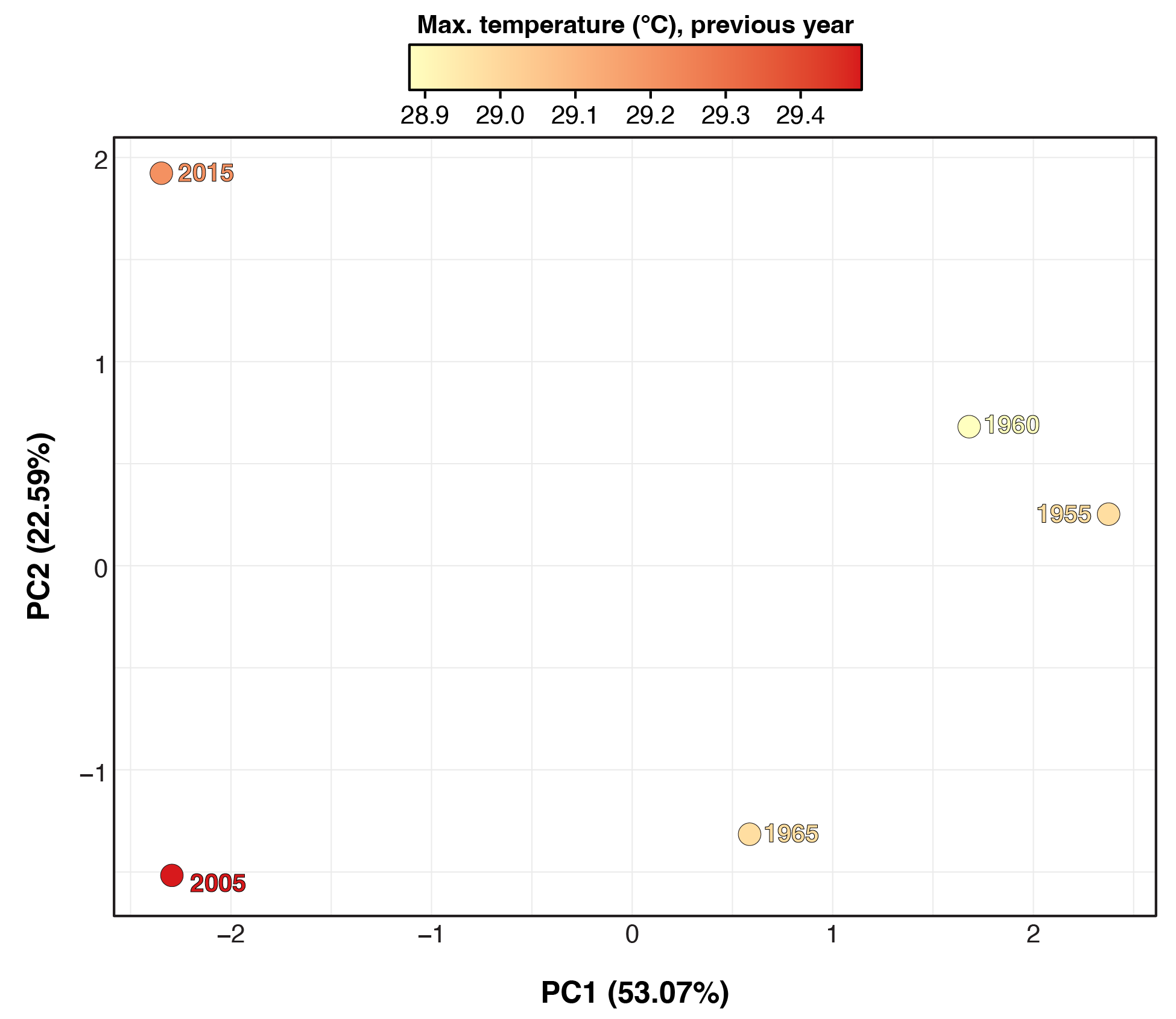

**Supplementary Figure 8.** Principal Component Analysis (PCA) calculated from the relative abundance of the 36 MAGs. Each data point corresponds to a subsample (i.e., timepoint) and its color to the maximum annual temperature of the previous year following the legend on top.

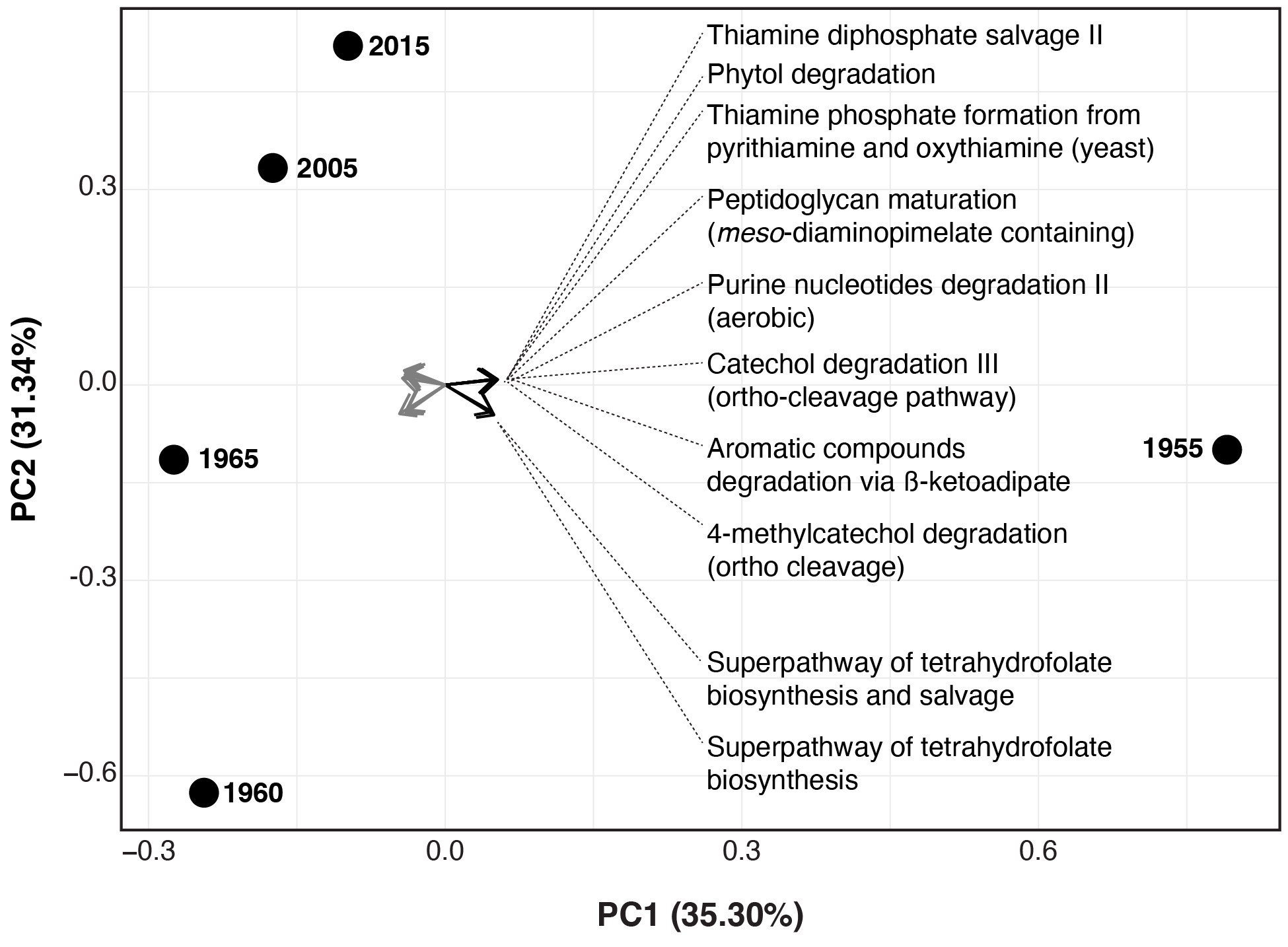

**Supplementary Figure 9.** Loading plot of the PCA in Figure 4b showing the top 10 pathways contributing to PC1 in both directions (positive and negative). Labels for those contributing to the PC1 positive direction are shown, for detail on those contributing to the negative direction see Supplementary Table 8.

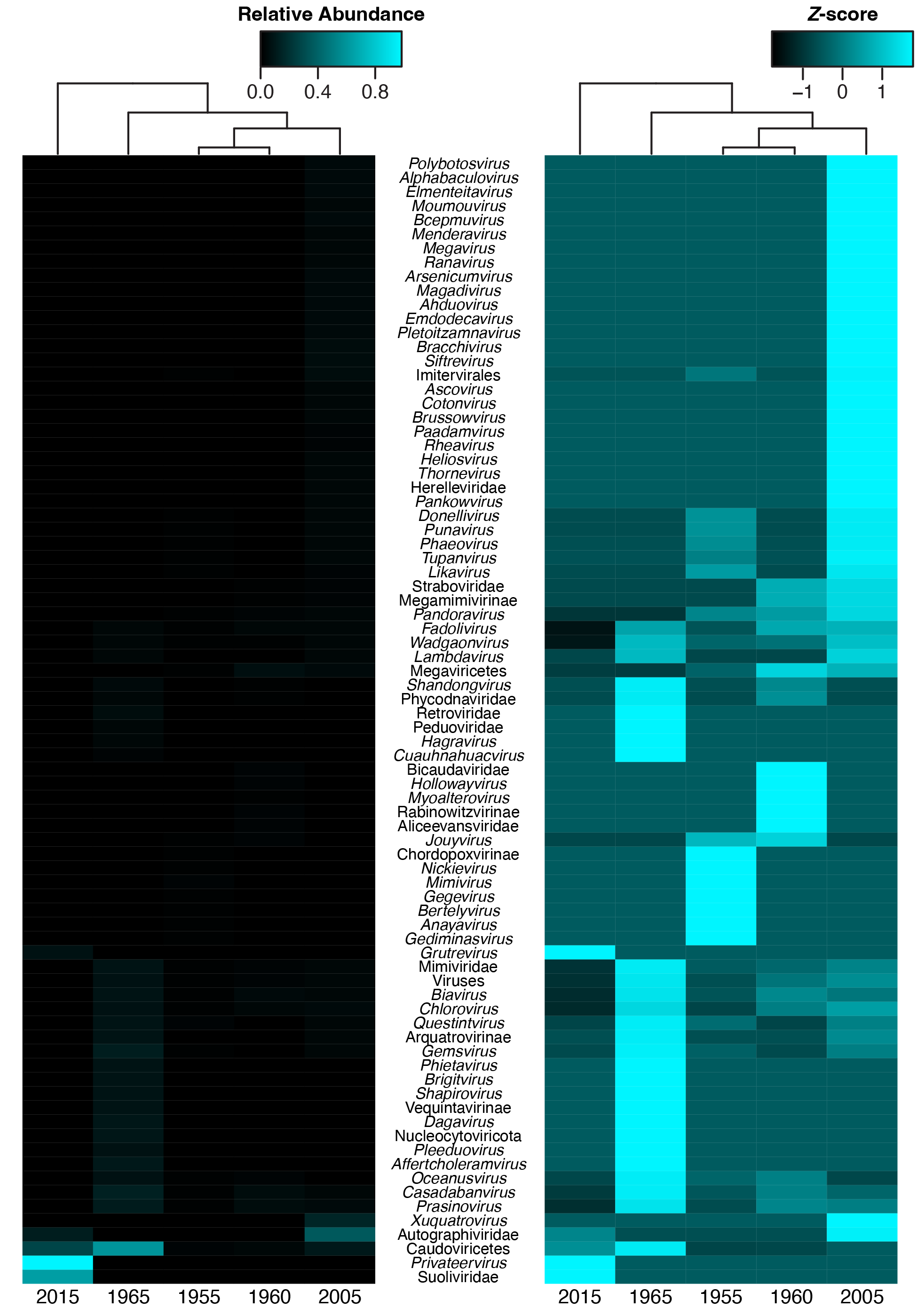

**Supplementary Figure 10.** Heatmaps of the Relative Abundance (left), and its *Z*-transformed values (right), of virus sequences identified using PhaBox in assembled MAGs (see *Experimental procedures*).

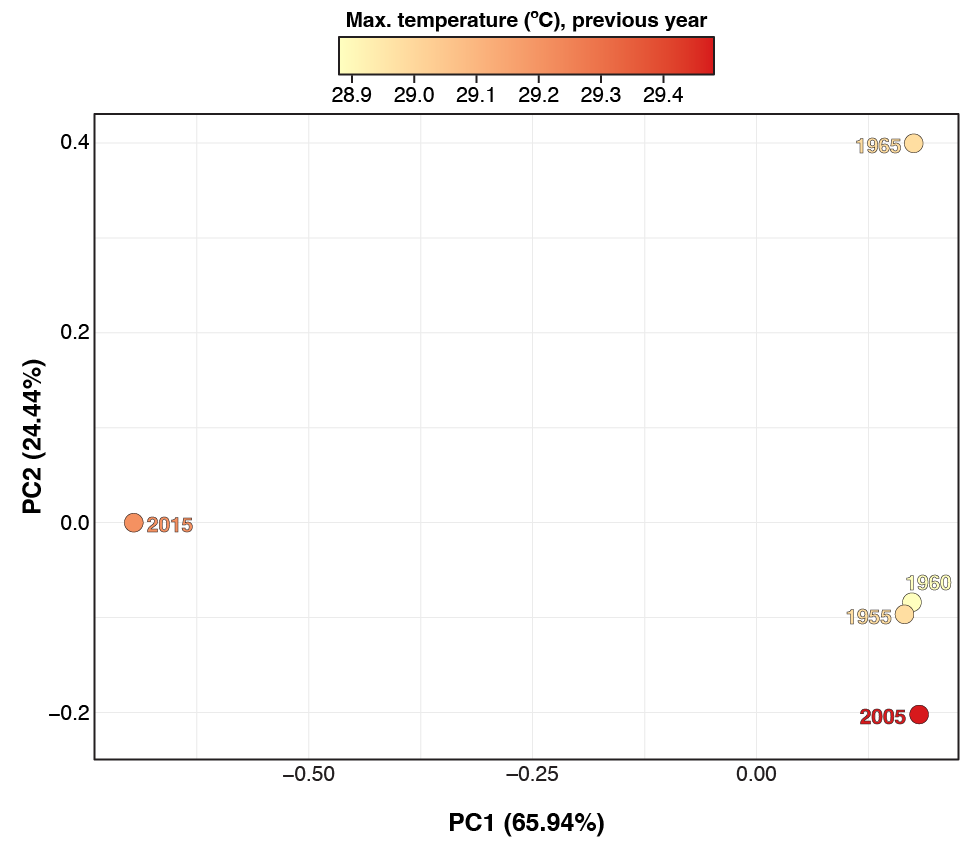

**Supplementary Figure 11.** Principal Component Analysis (PCA) calculated from the relative abundance of the PhaBox predicted viral sequences. Each data point corresponds to a subsample (i.e., timepoint) and its color to the maximum annual temperature of the previous year following the legend on top.

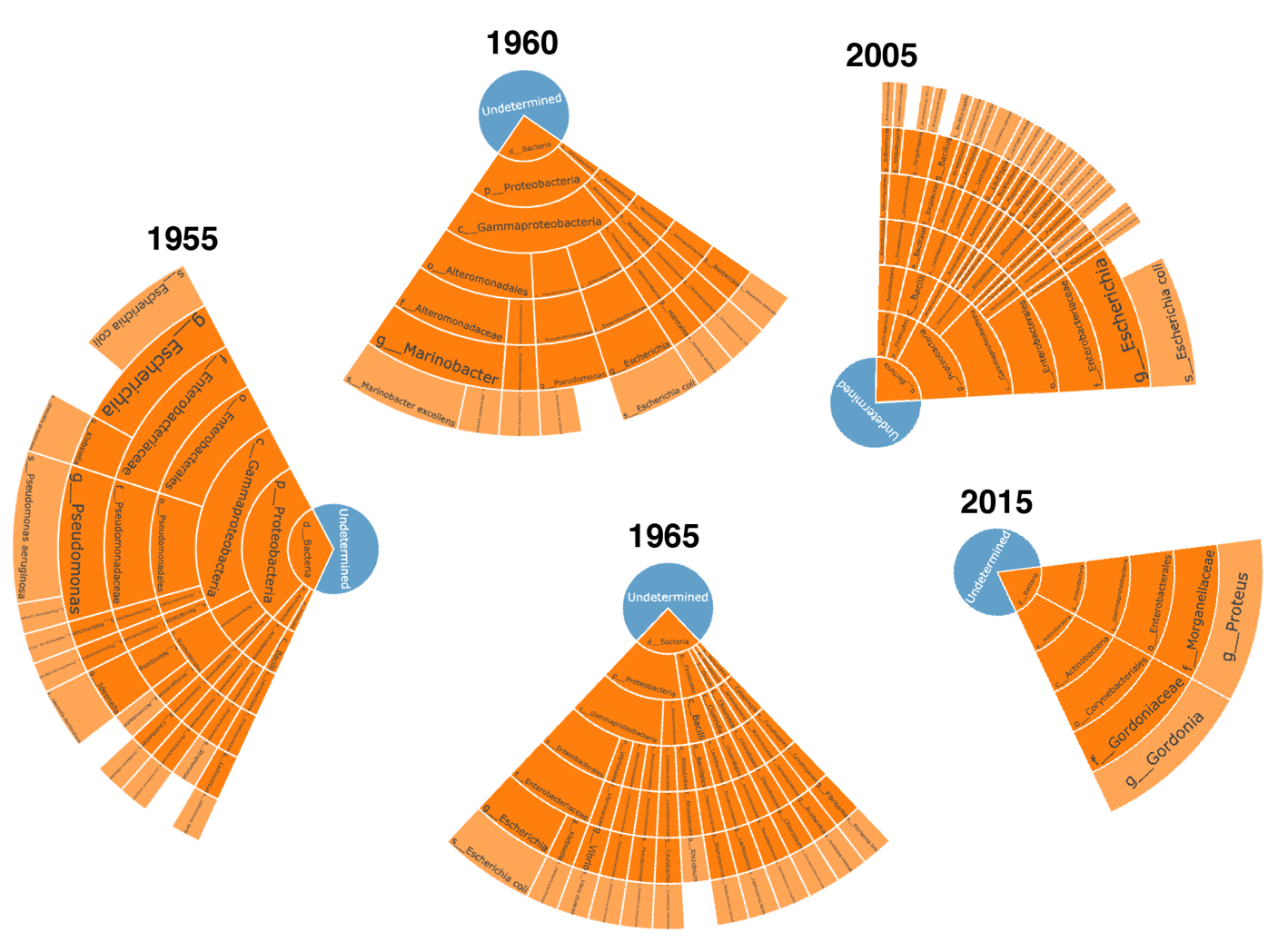

**Supplementary Figure 12.** Most abundantly predicted phage hosts by PhaBox.
